# Digital twins and hybrid modelling for simulation of physiological variables and stroke risk

**DOI:** 10.1101/2022.03.25.485803

**Authors:** Tilda Herrgårdh, Elizabeth Hunter, Kajsa Tunedal, Håkan Örman, Julia Amann, Francisco Abad Navarro, Catalina Martinez-Costa, John D. Kelleher, Gunnar Cedersund

**Author notes:** Contributed equally (Tilda Herrgårdh, Elizabeth Hunter).

## Abstract

One of the more interesting ideas for achieving personalized, preventive, and participatory medicine is the concept of a digital twin. A digital twin is a personalized computer model of a patient. So far, digital twins have been constructed using either mechanistic models, which can simulate the trajectory of physiological and biochemical processes in a person, or using machine learning models, which for example can be used to estimate the risk of having a stroke given a cross-section profile at a given timepoint. These two modelling approaches have complementary strengths which can be combined into a hybrid model. However, even though hybrid modelling combining mechanistic modelling and machine learning have been proposed, there are few, if any, real examples of hybrid digital twins available. We now present such a hybrid model for the simulation of ischemic stroke. On the mechanistic side, we develop a new model for blood pressure and integrate this with an existing multi-level and multi-timescale model for the development of type 2 diabetes. This mechanistic model can simulate the evolution of known physiological risk factors (such as weight, diabetes development, and blood pressure) through time, under different intervention scenarios, involving a change in diet, exercise, and certain medications. These forecast trajectories of the physiological risk factors are then used by a machine learning model to calculate the 5-year risk of stroke, which thus also can be calculated for each timepoint in the simulated scenarios. We discuss and illustrate practical issues with clinical implementation, such as data gathering and harmonization. By improving patients’ understanding of their body and health, the digital twin can serve as a valuable tool for patient education and as a conversation aid during the clinical encounter. As such, it can facilitate shared decision-making, promote behavior change towards a healthy lifestyle, and improve adherence to prescribed medications.

## Introduction

Healthcare is currently moving towards P4 medicine - personalized, predictive, preventive, and participatory medicine (1). This development away from reactive medicine, towards P4 medicine, is driven and facilitated by the increase in chronic diseases (2), by a general increase in mechanistic understanding behind disease development, and by technological advancements (3,4).

One already existing clinical practice, which has the potential to include all P4 principles, is the preventive health dialogue (Figure 1A). During such a health conversation, an individual and a specialized nurse discuss different health aspects regarding the individual’s current lifestyle and health status. The discussions cover the current and projected risk of getting different lifestyle related diseases, such as a stroke. The discussions also cover what an individual can do to decrease that risk (Figure 1A and B). With this as a basis, the individual decides which lifestyle changes to do, and is also responsible for maintaining those lifestyle changes (Figure 1C).

**Figure 1:**
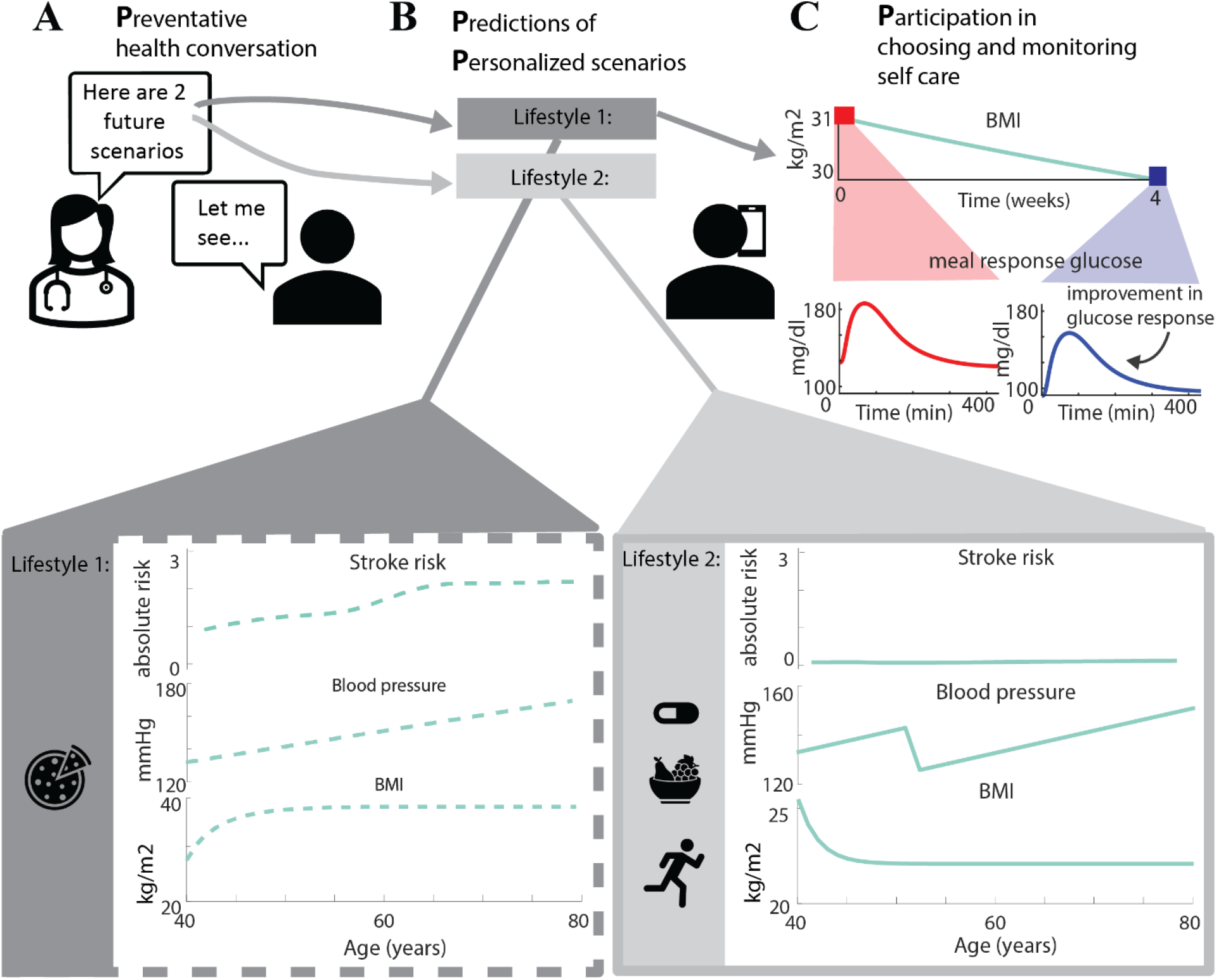
Vision for P4 medicine - preventive, predictive, personalized, and participatory medicine. A) during preventive health dialogues, different future scenarios are presented and discussed. Here, two examples of such scenarios are shown - B) Lifestyle 1: The person does not take any blood pressure medication. The person continues the unhealthy lifestyle, thereby increasing BMI, and eventually developing diabetes. The risk increases during the simulated 40 years. Lifestyle 2: The person starts to lose weight at 40, consequently decreasing BMI, and starts to take blood pressure medication when turning 50. The risk of stroke does not increase much during the 40 years. C) The person participates in his/her own care by choosing what lifestyle changes to make and by continuously monitoring the progress and results of the lifestyle changes. Here, this monitoring is exemplified by looking at how weight loss intervention results in improved meal glucose response. By continuously comparing such predictions with data, the person can continuously deepen their understanding of their own body, and hopefully better maintain motivation.

While the existing health dialogue already features some aspects of the P4 principles, these principles could be further strengthened in various ways. Firstly, today, the health dialogue does not make use of advanced predictions using personalized computer models. In Figure 1B, such predictions show the effect of two different lifestyles on the risk of a stroke, by simulating physiological variables such as blood pressure and BMI. Secondly, another possibility with computer models is to zoom in on things happening on shorter timescales and other biological levels, using multi-level and multi-timescale models (Figure 1C). By simulating short-term changes, one can compare those predicted changes with measured outcomes during different time points. If those predictions turn out to be correct, it has the potential to increase faith in the long-term predictions, give continuous feedback on progress, and thereby increase motivation. In Figure 1C, this potential is exemplified with a simulated long-term decrease in BMI, which is associated with changes in short-term meal response. One technology that could contribute to all these possibilities is digital twins.

A digital twin of an individual is a personalized computer model of that individual, and it has been suggested that such twins can be used for prevention of cardiometabolic diseases such as diabetes and stroke (5–7). Digital twins can be described by black box phenomenological models or by mechanistic physiology-based models. Phenomenological black box models are suited for making predictions and statistical assessments, such as the risk of a stroke. In contrast, mechanistic models describe physiology and biochemistry using mechanistic data, typically time series. These mechanistic models are first developed using varying amounts of *in vitro, in vivo,* and clinical population data, to be able to describe the physiological and biochemical processes (8). Then, these models are individualized using data from the particular individual, to be able to make personalized predictions. Finally, a twin can be either online, if it is continuously updated, or offline, if it is occasionally updated (9)

One area where digital twins are especially suited is for preventative care of cardiovascular diseases such as stroke. These diseases are at least in part preventable with life-style changes and early medical interventions, and have a complex pathophysiology. The complexity of cardiovascular diseases comprises several medical and environmental factors, and involves multiple organs, timescales, and control mechanisms (10). This complexity makes it difficult to make individual predictions without help from advanced analytical tools such as digital twins. Capturing the complexity with a digital twin could potentially help physicians and patients choose patient-specific preventive measures, by first testing the different measures on the digital twin and thus identifying the one with the most promising effects for that individual. Despite this potential, digital twins have only rarely been used in healthcare, partly because the models are not yet ready for usage.

One of the reasons that the models are not ready for usage is that the development of these models has relied upon either of two different modelling approaches: mechanistic or black box machine learning models. Mechanistic models describe the underlying physiological processes, leading to changes over time (Figure 2A). Specifically, they can take lifestyle variables (e.g. energy expenditure) and personal parameters (e.g. age) as input, and can then be used to simulate the evolution of biomarkers and their covariation with other biomarkers. These kinds of models can then be used to simulate different scenarios (e.g. different lifestyles) and compare them with each other, on both the long term-level and on short-term effects. Since this approach directly uses prior physiological understanding to develop the models, the resulting models can give comprehensive physiological explanations for their predictions. Mechanistic models are primarily used for improved mechanistic understanding or for predicting the dynamics of physiological variables. Machine learning models, on the other hand, can map between inputs and outputs, such as risk factors and risk scores, without needing any information about the physiological mechanisms in that mapping. These models usually do not learn anything new about the mechanisms of the physiological system that produced the predicted output. Machine learning has within health care been successful in e.g. classifying images (11) and calculating risk scores (12–15). Both modelling approaches have different needs regarding data: machine learning models need a lot of data (1000s of subjects), preferably from different sources/demographics to ensure generalizability, and mechanistic models need time-series data for the response to some perturbation of a system, e.g. a meal or a medication.

**Figure 2:**
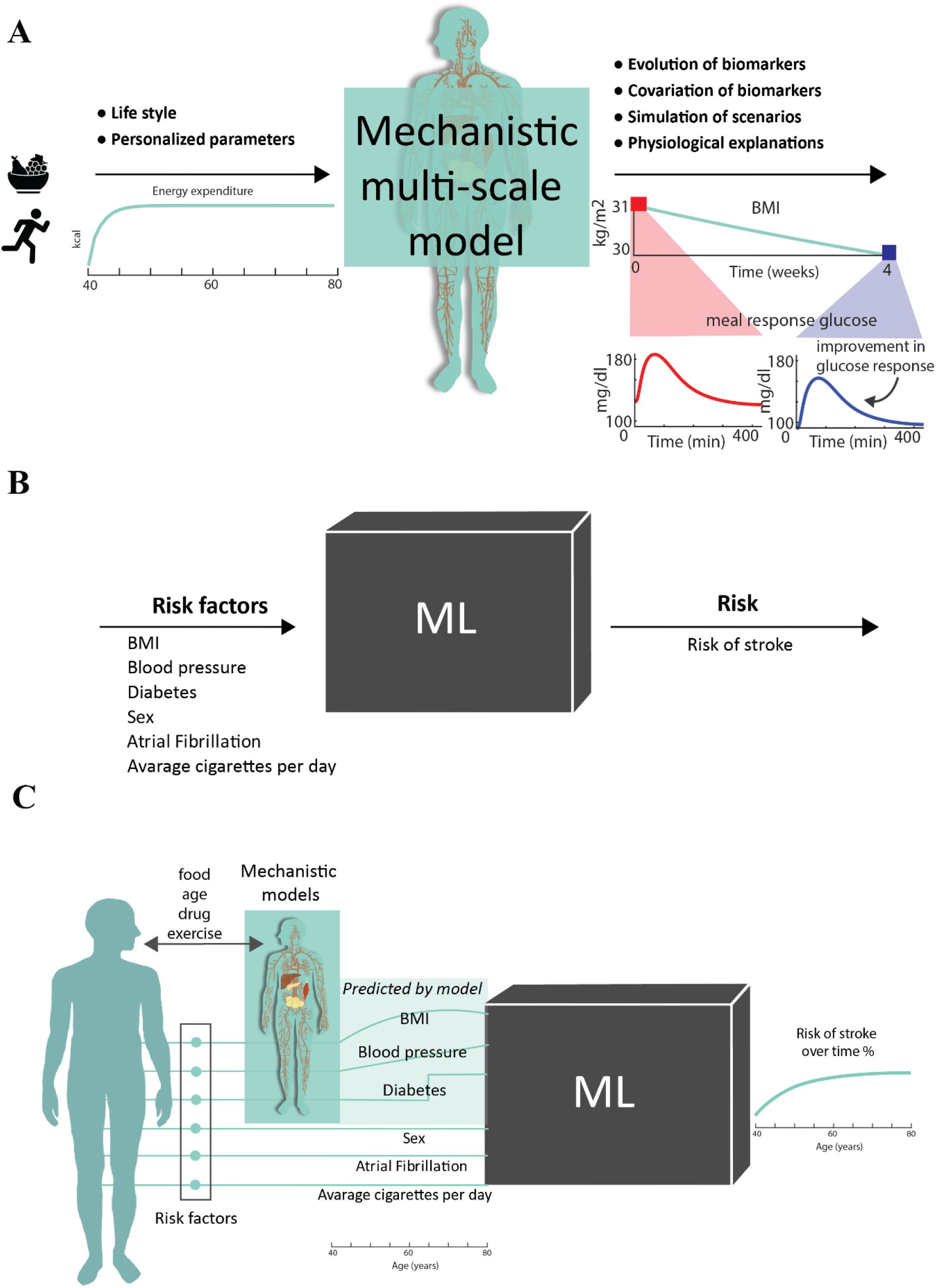
Two different modelling approaches, mechanistic and machine learning can be used on their own and in a combined hybrid scheme. A) Mechanistic models can make use of personal parameters and lifestyle data to simulate the evolution of different biomarkers (e.g. BMI and glucose meal response). As such, mechanistic models can be used to simulate different scenarios (e.g. different lifestyles or drugs) and give physiological explanations of these predictions. In this figure, an increase in energy expenditure is used as input, and the model is used to simulate BMI over 80 years, as well as meal glucose levels at different time points during these 80 years. B) Machine learning models can use different risk factors as input and give an estimated risk score as output. These models are so called black box models, i.e. one cannot look at their model structure and discern any physiological understanding. C) The complementary benefits of the two different modelling approaches can be utilized in a hybrid scheme. Here we suggest and present such a scheme. First, data from an individual are used to personalize the mechanistic model. Second, lifestyle inputs, such as food, age, drug, exercise, are used to make simulations of relevant biomarkers over time. Finally, these simulations, together with other risk factors not simulated by the mechanistic models, are then used to simulate the time-varying risk of stroke.

Since the two modelling approaches have complementary abilities, a combination of the two has sometimes been proposed (10,16–19), even though few examples exist. Such combinations have for instance been used to improve MRI classification (20–22), for predicting tumor growth and density (23), and for predicting phenotypes based on genotypes (24). Hybrid schemas, where ML is used to formulate the mechanistic models, or where mechanistic models are used to constrain ML models, have also been proposed and developed (17,25). For example, ML techniques have been used to train and develop a mechanistic network model that were then used to predict cell responses to combinations of perturbations (e.g. drugs) (26). Within the field of cardiovascular diseases specifically, a hybrid model that combines a discrete model with an agent-based model has been used to simulate treatment procedures of heart failure and comparing both health outcomes and financial aspects (27). However, despite these many suggestions and first examples, and despite the importance of developing digital twins for stroke, no hybrid model underlying digital twins for stroke has yet been developed.

Herein, we present a digital twin for stroke prediction, combining multi-level and multi-timescale mechanistic models with a machine learning model that calculates the 5-year risk of stroke, and show how it can be used in health care. The mechanistic models combine a model for the progression of type 2 diabetes with a new model of how blood pressure changes with age and medication (Figure 3A). We show how this mechanistic model can simulate how different lifestyle choices lead to different scenarios for the evolution of both the overall risk of stroke, and for known physiological risk factors, such as weight, diabetes development, and blood pressure (Figures 9-11). Finally, we discuss outline solutions to practical issues regarding implementation, such as data harmonization, and ethics.

**Figure 3:**
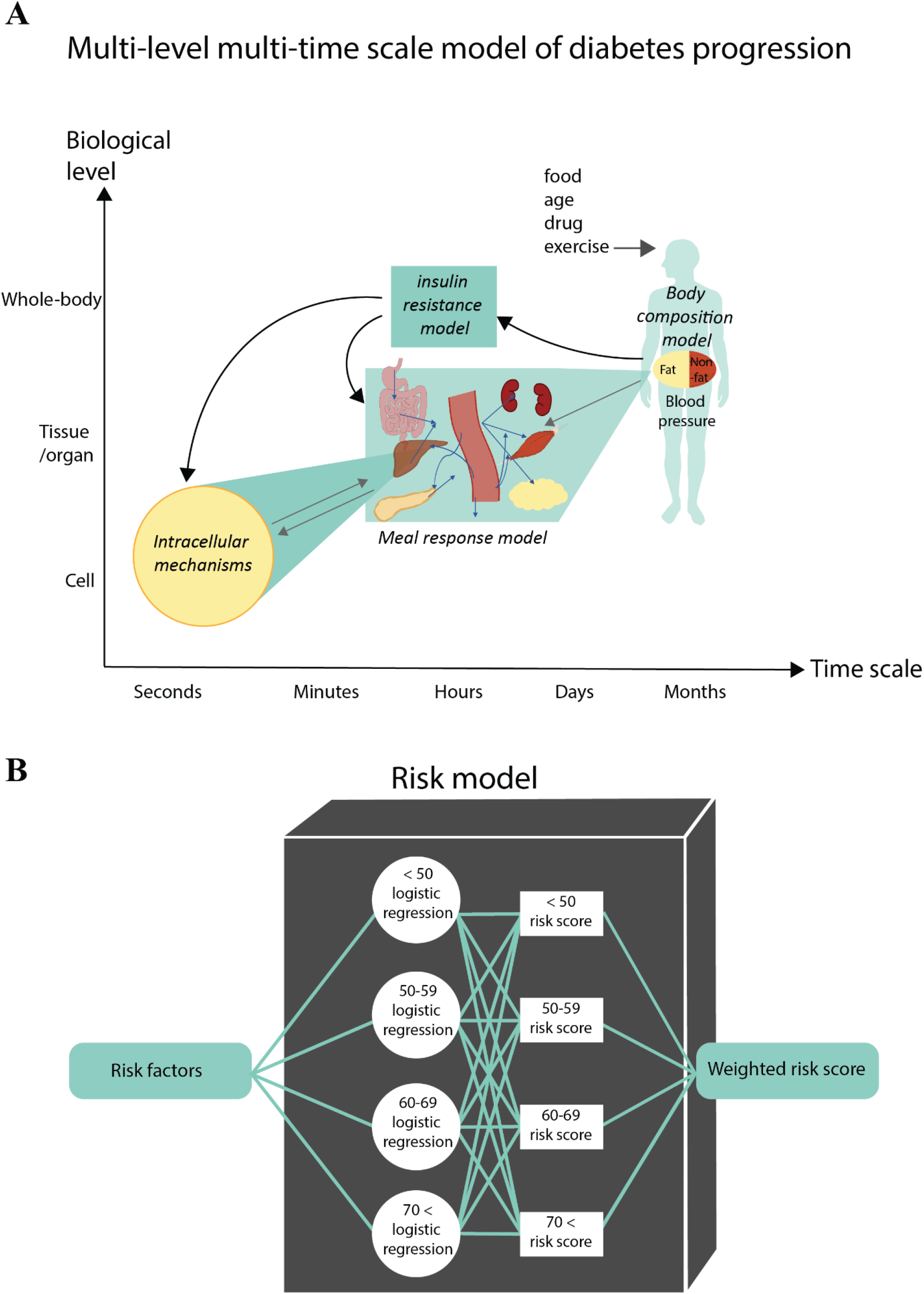
The models included in the herein presented hybrid model. A) The mechanistic part consists of three different biological levels: whole-body, organ/tissue, and cell level. The models can also simulate dynamics on different time scales: on seconds and minutes up to days and months, even years. B) The risk model consists of an ensemble of 4 different logistic regression models for 4 different age groups, taking in the same risk factors as input. The final risk score is a weighted sum of the risk scores from each of the models.

**Figure 4:**
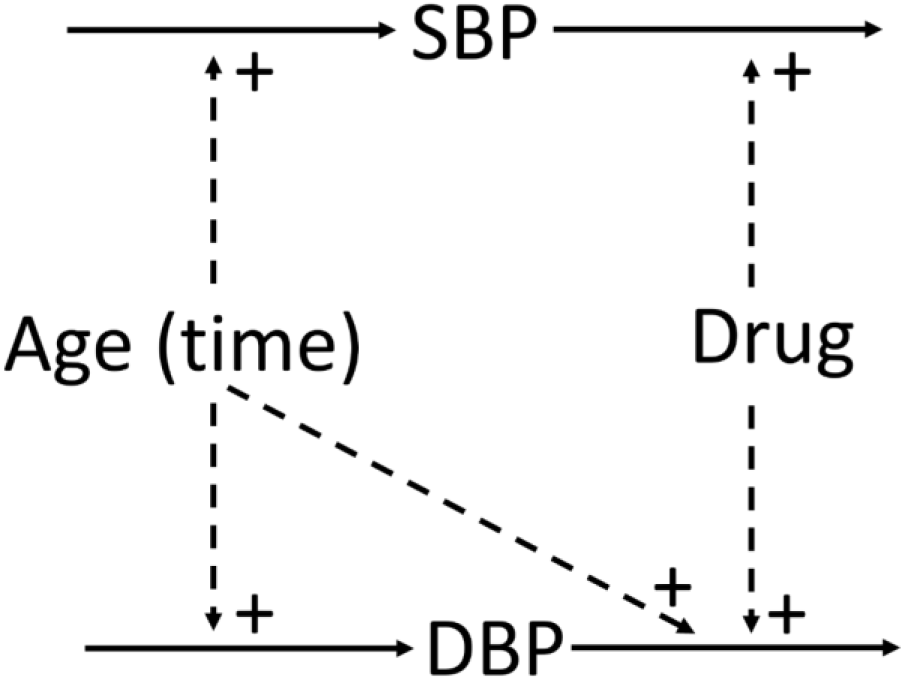
Overview of the blood pressure model. SBP is systolic blood pressure, DBP is diastolic blood pressure, and both are affected by both age and by the drug.

## 2 Method

### 2.1 Mechanistic models

The mechanistic models are built up by ordinary differential equations (ODEs) in the standard form:

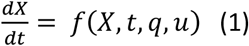

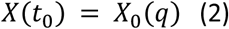

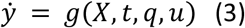

where *X* is a vector of state variables, *f* and *g* are non-linear smooth functions, *g* a vector of model parameters, u an input signal, *X*(*t*_0_) initial conditions, and 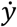 is the simulated model output.

The complete interconnected model consists of four different sub-models describing metabolic control at three different levels: cell, organ/tissue, and whole-body (Figure 3A). All of the equations are given in the Supplementary material (which is based on (28,29)) both as equations and as simulation files. The initial values were obtained through steady state simulation, set to match the scenarios, or kept the same as in original articles. All initial values used in the simulations can be found in the Supplementary material. The fit of the 4 insulin resistance parameters were adjusted by hand, to get a diabetic behavior of relevant variables, corresponding to that of the diabetes parameters in (30). The simulation of the mechanistic model, and numerical optimization of model parameters, were carried out using MATLAB 2018b and the SBtoolbox2 package.

### 2.2. Parameter estimation in the blood pressure model

The parameter estimation, done only for the new blood pressure model, was done using the traditional weighted least square cost function:

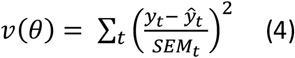

where *y_t_* is the measured data at timepoint *t, ŷ_t_* the simulation value at time point *t*, and *SEM_t_* is the standard error of mean at time point *t*. This cost function is then minimized over the free parameters. Only the parameters of the new blood pressure model were fit to data, and there is no cross-talk between the blood pressure model and the metabolic models. Two datasets were used: Framingham and VALUE. For the Framingham data, the blood pressure parameters were optimized to minimize the cost using two MATLAB functions, first particleswarm and then simulannealbnd. For the VALUE data, the model parameters were manually adjusted to fit to data. Since no standard deviation was given in the Framingham data, the standard deviation was approximated to 3 in the SBP and 2 in the DBP when using a *χ*2-test for the training, and standard deviations of 4 for SBP and 2 for DBP in the cross-validation.

### 2.3 χ2-test

The parameter estimation was evaluated using a χ2-test of the size of the residuals. The null hypothesis for this test is that the experimental data have been generated by the model with additive and normally distributed noise. The test statistic value that is compared with the cost function is given by the inverse cumulative density function,

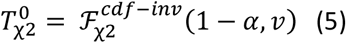

where 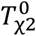 is the test statistic, 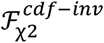 is the inverse cumulative density function, α is the significance level (*α* = 0.05 was used), and *v* is the degrees of freedom, which was equal to the number of data points in the training dataset. If the model cost is larger than the χ2-threshold, the model is rejected.

### 2.4 Blood pressure model

The new blood pressure model consists of 2 states:

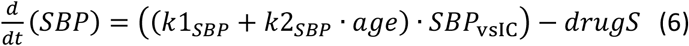

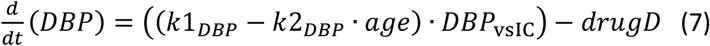

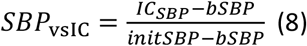

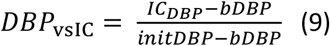

Where *SBP* and *DBP* are blood pressures, *drugS* and *drugD* are the drug effects, *k*1_SBP_, *k*2_SBP_, *k*1_DBP_, *k*2_DBP_, *k*1_DBP_, and *k*2_DBP_ are parameters, *SBP*_vsIC_ and *DBP*_vsIC_ are patient-specific constants that depend on the initial blood pressure values at age 30, parameters *bDBP* and *bSBP* are parameters that determine the difference in the age-related increase in blood pressure between different patients with different initial blood pressure values, and *initSBP* and *initDBP* are constants set to the initial values in the non-hypertensive group 2 in the Framingham data.

### 2.5 ML risk model

The used machine learning risk model is based on a set of four independent logistic regression models that are designed to predict the 5-year risk of initial ischemic stroke for an individual, belonging to four different age groups, as already presented in (31). There is a logistic regression model for each of the following four age groups: under 50, 50-59, 60-69 and over 70. Each of the four age-specific models has the same risk factors as predictors. The risk factor factors included in the model are: sex, systolic blood pressure (SBP), diastolic blood pressure (DBP), BMI, average cigarettes smoked per day, atrial fibrillation and diabetes. To determine an individual’s risk score, their risk factors are used as input into the appropriate model for the individual’s age. To use the age models in the hybrid modelling approach, we have adapted the independent set of logistic regression models into an ensemble model as described below.

Age-specific models have been shown to capture the non-proportionality of risk factors by age and are better calibrated to the younger and older populations compared to a model that is trained on all ages with age included as a risk factor. Thus, age-specific models are particularly suited to this hybrid modelling approach that aims to calculate the risk based on simulated scenarios across time, as they will identify and use the risk factors that are most important to the individual’s stroke risk at a given time point (31).

However, there are some disadvantages associated with the use of independent age-specific models for the hybrid model. In particular, when the age switches from one age-group to another, there may be a discontinuous jump in the risk, since one switches from one model to another. To avoid this problem, instead of assigning an individual to a given age group and basing their stroke risk solely on the prediction of the age specific model corresponding to their age group, we calculate their risk in each age group in parallel and then take a weighted average of these risks. In other words, we convert the set of age-specific models from a set of independent models to an ensemble, with the predictions from each of the models being integrated to generate a single overall risk for an individual. To weigh the stroke risk score for an individual returned by each age-group specific model, we use the distance from the individual’s age to the corresponding age group of the model that returned the risk score. The function used to calculate the distance *d* is given in Equation 10 which returns the square root of the sum of the difference between an individual’s *Age* and the minimum value in an *AgeGroup* squared with the difference between an individual’s age and the maximum value in an age group squared.

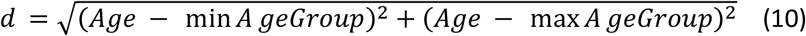

Where *Age* is the age, and *AgeGroup* is the age group. Within an age group, the smallest value for the distance will be for the age in the middle of the age group and the largest distance will be at the maximum or minimum age. For example, in the 50 to 59 age group, an individual aged 54 or 55 would have the smallest distance to the group while an individual aged 50 or 59 would have the largest. For those not in the age group, the ages closest to the group will have the smallest distance. Someone aged 60 would have a shorter distance to the 50 to 59 age group than someone aged 70. For an individual, the distance to each age group is calculated and then these distances are used to weigh risk scores from corresponding age group specific models and the weighted sum of the risk scores is the individuals total risk score. If we take the example of a 57-year-old person, they will have four distance parameters, d_50_, d_5059_, d_6069_ and d_70_, and four weights w_50_, w_5059_, w_6069_, w_70_. These distances would be calculated as follows:

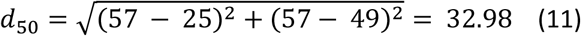

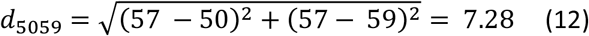

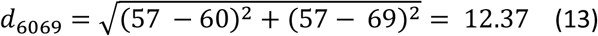

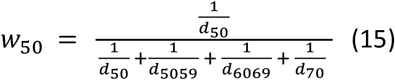

The weights for each risk model are created using inverse distance weights and are defined below:

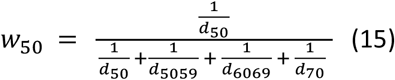

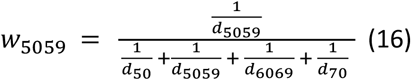

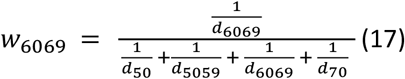

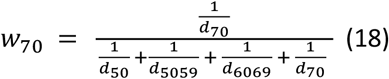

For the 57 year old, if we plug the distances found in Equations 11 - 14 we get the following weights: w_50_ = 0.11, w_5059_ = 0.50, w_6069_ = 0.30, and w_70_ = 0.09.

To calculate the final risk score we then use the following formula where the variables R_50_, R_5059_, R_6069_, R_70_ are the risk scores calculated by each of the four independent age group specific models:

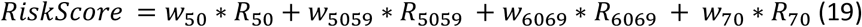

For the 57 year old, we see that the risk score from the 50-59 age model is weighted the highest followed by the risk score from the 60-69 age model; then the under 50 model and finally the over 70 model.

### 2.6 Code availability

*All code will become available, when this manuscript has been accepted in a journal.*

## 3. Results

### 3.1. Development and validation of the new mechanistic blood pressure model

The blood pressure model describes the effect of aging on systolic and diastolic brachial blood pressure. The model of aging was trained on data from the Framingham Heart Study (32), including four groups with different SBP at the end of the study (Figure 5). The model passes a χ2-test for all 4 groups (V(θ)=32.5 < 111= χ^2^_cum,inv_(88,0.05)). A model validation was performed on the same data by leaving out one group during model training and then use the left-out group as validation data (Figure 6). The cross-validation in the Framingham data also passes a χ^2^-test (V(θ)= 3.03, 0.97, 17,2, 26.5<33.9 = χ^2^_cum,inv_(22,0.05)).

**Figure 5:**
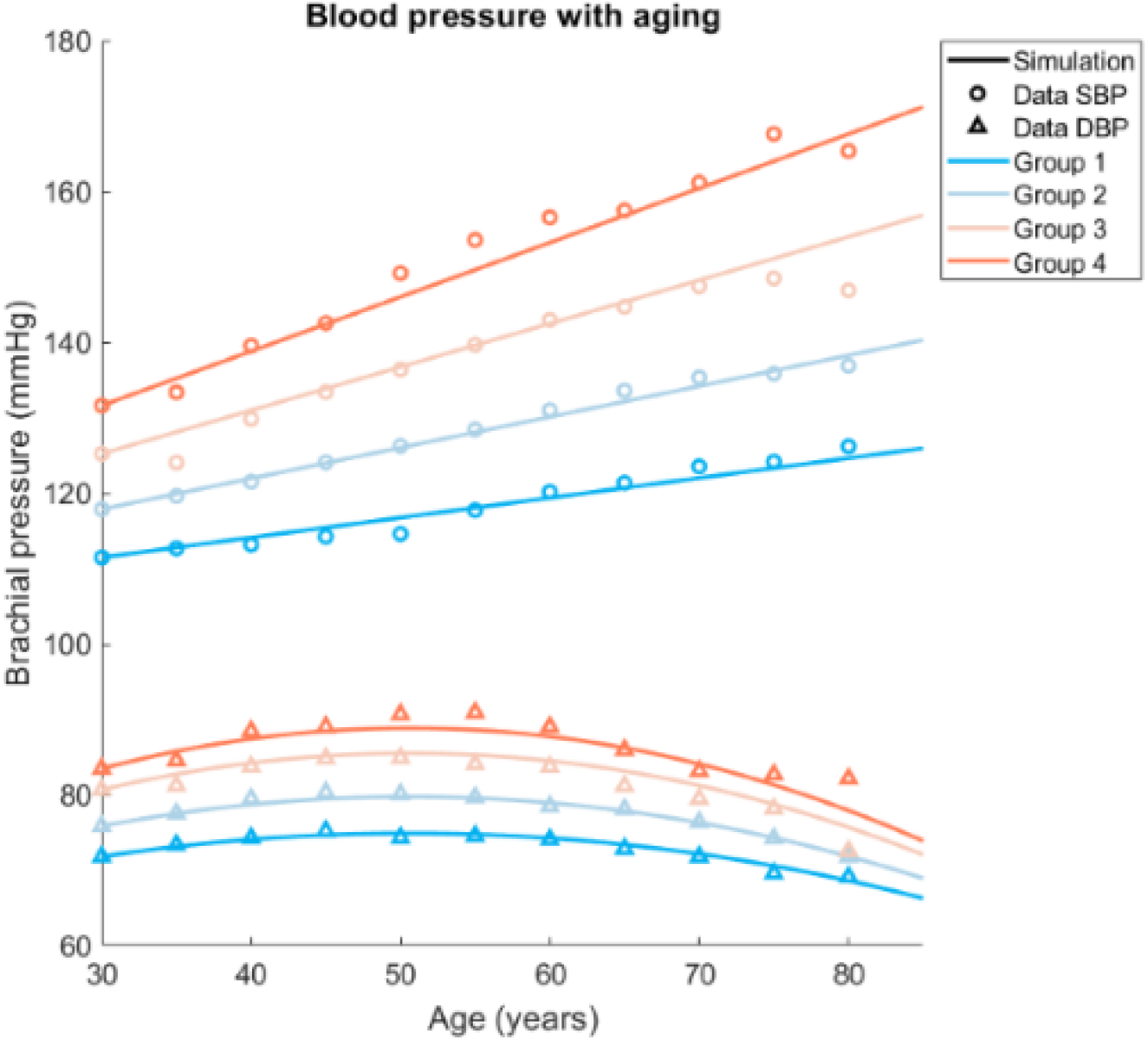
Fit to data of aging from Framingham Heart Study.

**Figure 6:**
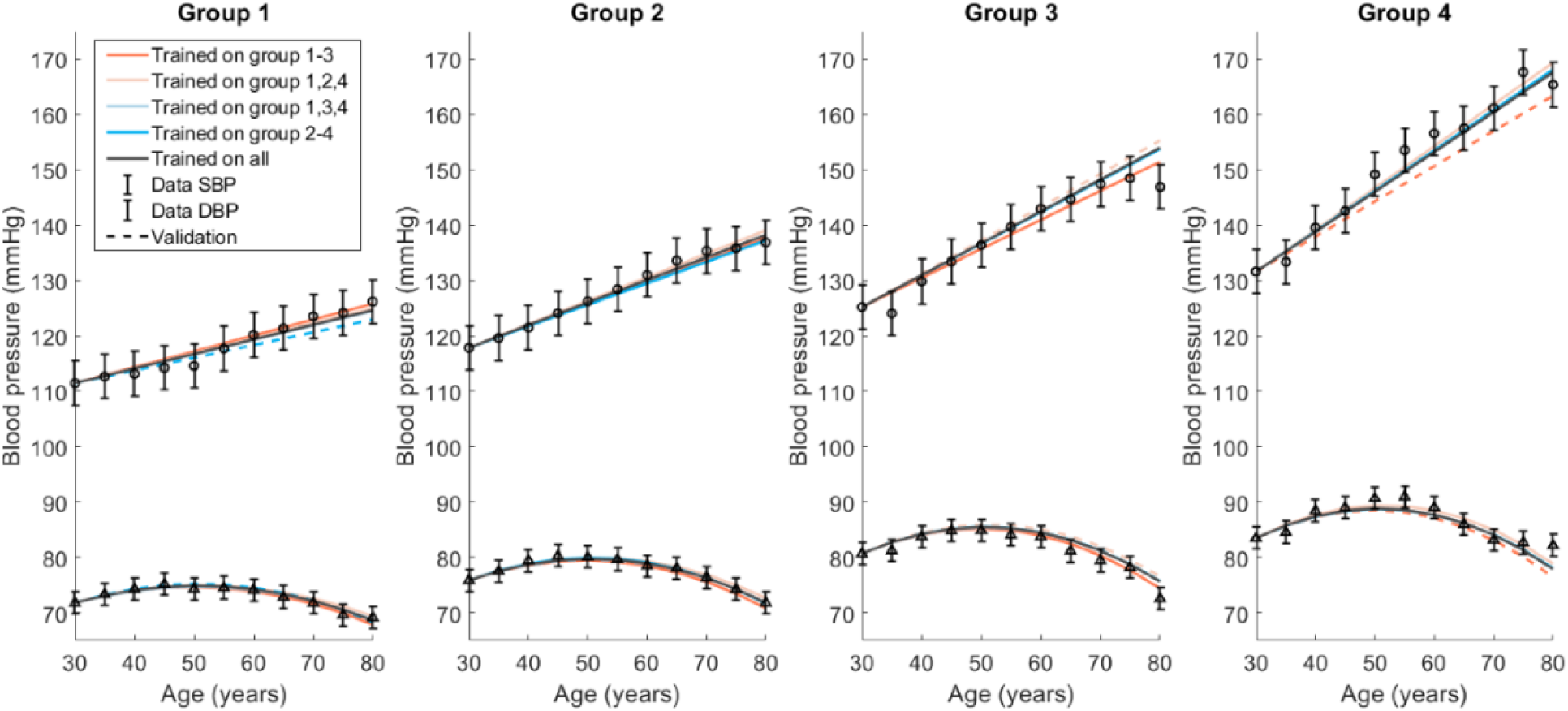
Cross-validation on data from the Framingham Heart Study.

A drug mechanism was added to the model to describe the decline in blood pressure due to anti-hypertensive drugs (Figure 7). The combined long-term model of blood pressure was then trained on data from an angiotensin receptor blocker (ARB)-based treatment from the VALUE trial, where treatment started at the age of 67.2+-8.2 (33). The fit of the trained model to the drug-treatment data passes a χ^2^-test (V(θ)=0.8 < 26.3 = χ^2^_cum,inv_(16,0.05)).

**Figure 7:**
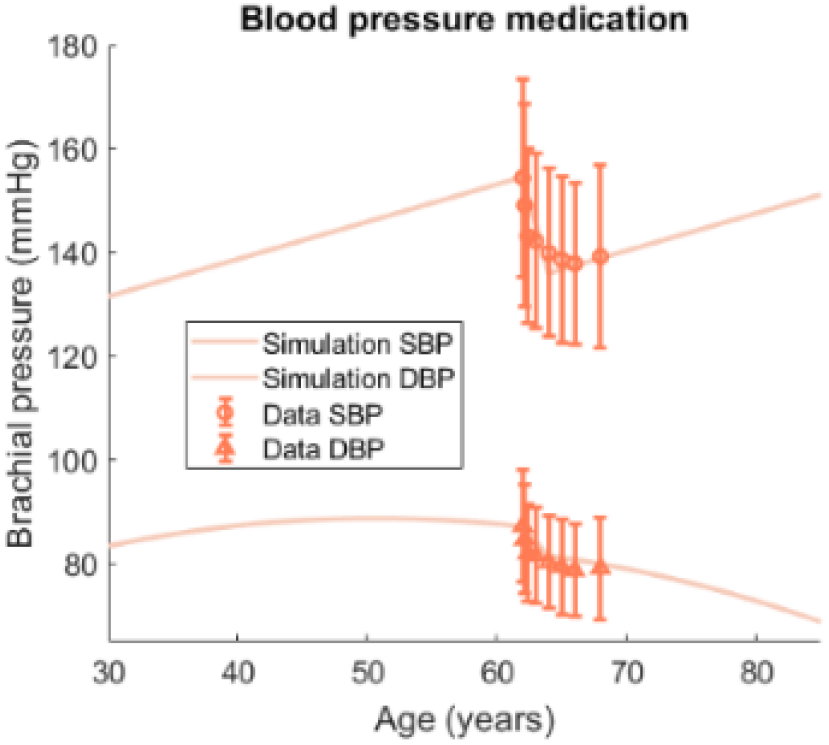
Fit to data of Valsartan-based treatment, where the final trained model was used to simulate an ARB-based treatment starting at the age of 45 compared to no treatment.

**Figure 8:**
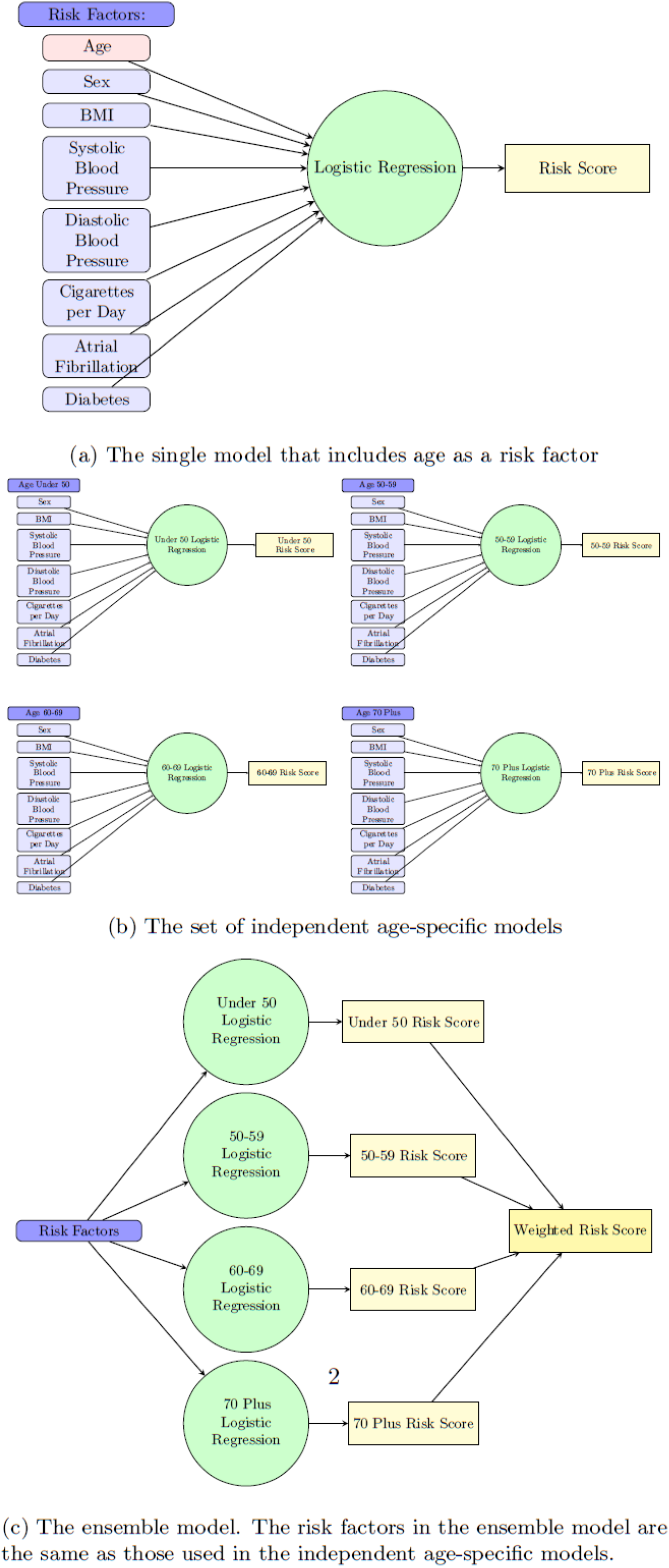
The ensemble model.

The VALUE trial data were also used to choose the timepoint for start of drug treatment used in the hybrid simulation of blood pressure (Figure 10). This timepoint was chosen as the timepoint where the group from the Framingham-data with the highest SBP had the closest SBP to the mean baseline value from the VALUE trial and was found at t=62 years. This choice of timepoint was based on the fact that the person in the simulated scenario is within the age span of the subjects from the VALUE trial (67.2+8.2). The reduction in blood pressure was assumed to be the same for all values of the blood pressure at the start of treatment. The simulation of normal aging used is the same as was fitted to all four groups from the Framingham study.

### 3.2. Validation of the ensemble-based risk model

The independent age-specific models, without the ensemble approach, were evaluated in (31). However, since we do a new ensemble-based combination of these age-specific models, it is necessary to look at the evaluation metrics of the ensemble model. We look at discrimination and calibration for the model as a whole and for each of the age groups using an 80:20 train:test split, where we use 80% of each age group data set to train the corresponding age group specific model and then combine the left over 20% of data from each age group to test the weighted model. Looking at both discrimination and calibration is important to fully understand the performance of the model. While the discrimination metrics provide a measure of the predictive capabilities of the model, the calibration measures the difference between the estimated and true risk, which can help to identify if the model does not predict well for certain groups (34). For discrimination we look at the F1 measure, the AUC, and the accuracy of the model. The F1 measure is the harmonic mean of how often a model makes a positive prediction that is true (precision) and the true positive rate (recall). The AUC is the area under the receiver operating characteristic (ROC) curve which plots the true positive rate against the false positive rate as the prediction threshold of the model moves between 0 and 1. Accuracy is the portion of predictions that are correct (35). To look at calibration, we use the Spiegelhalter’s Z-test, which separates the calibration aspects out of the Brier Score (the mean squared error between the outcome and the estimated probabilities), and the Hosmer-Lemeshow test, which examines goodness of fit. If the p-value is significant in either of these two tests, the model is assumed to be not well calibrated.

Using both the discrimination and calibration metrics described above, we compare the ensemble of age-specific models with two models from (31): a set of independent age-specific models (i.e., where the predicted risk for an individual is based solely on the prediction of the age-group specific model that covers their age), and a single model that includes age as a risk factor and that is trained on data from all the age groups. Note that the set of independent age-specific models are, in fact, the component models that are used to construct our ensemble model. In other words, the comparison between these two models directly assesses the contribution of our function for weighting different risks. Figure 8 provides a visualization of the three stroke risk prediction models.

#### 3.2.1 Discrimination analysis of the stroke risk model

Table 1 shows the values for AUC, F1, and accuracy for the ensemble of age-specific models tested on the individual age groups, as well as the same measures for the set of independent age-specific models and the single model with age as an input. Comparing the results in Table 1, we see that the set of independent age-group specific models has the highest AUC in all categories and the single model with age as an explicit input (which is representative of a standard approach to stroke risk modeling) has the lowest AUC in all age categories. With respect to F1 the ensemble of the age-specific models has the best performance in all age categories except 60-69, where the independent age specific model has best performance. Finally, looking at accuracy, the independent age-specific models have the best accuracy in the younger age-groups (Under 50, and 50-59). However, in the older age groups the single model with age as an input has slightly better performance and the performance of the ensemble of age specific models either matches or exceeds the performance of the independent age-specific models. Overall, however, we conclude that our new ensemble-approach used to smooth out transitions between the age group model still produces a roughly equal discriminative ability.

**Table 1:** Discrimination Metrics

|  | AUC | F1 | Accuracy |
| --- | --- | --- | --- |
| **Under 50** |  |  |  |
| Ensemble of Age-Group Models | 0.57 | 0.37 | 0.66 |
| Independent Age-Group Models | 0.67 | 0.22 | 0.70 |
| Single Model | 0.52 | 0.31 | 0.64 |
| **50-59** |  |  |  |
| Ensemble of Age-Group Models | 0.66 | 0.45 | 0.68 |
| Independent Age-Group Models | 0.70 | 0.41 | 0.71 |
| Single Model | 0.64 | 0.43 | 0.66 |
| **60-69** |  |  |  |
| Ensemble of Age-Group Models | 0.71 | 0.41 | 0.68 |
| Independent Age-Group Models | 0.72 | 0.47 | 0.68 |
| Single Model | 0.69 | 0.37 | 0.69 |
| **70 Plus** |  |  |  |
| Ensemble of Age-Group Models | 0.69 | 0.49 | 0.72 |
| Independent Age-Group Models | 0.70 | 0.44 | 0.71 |
| Single Model | 0.69 | 0.49 | 0.73 |

#### 3.2.2 Calibration analysis of the new risk model

Table 2 shows the p-values for the two calibration tests for the ensemble of age-group-specific models, the independent age-group models, and the single model. From the table, we can see that in both the ensemble and independent age models, all age groups have p-values for the goodness of fit tests greater than 0.05, suggesting that these models are well-calibrated across all age groups. This is an improvement from the single-age risk model and suggests that the new ensemble approach does not alter the acceptable calibration property of the previous model (31).

**Table 2:** Calibration Metrics

|  | Hosmer-Lemeshow | Spiegelhalter's |
| --- | --- | --- |
| **Under 50** |  |  |
| Ensemble of Age-Group Models | 0.19 | 0.14 |
| Independent Age-Group Models | 0.23 | 0.34 |
| Single Model | 3.45e-5 | 0.002 |
| **50-59** |  |  |
| Ensemble of Age-Group Models | 0.17 | 0.40 |
| Independent Age-Group Models | 0.34 | 0.52 |
| Single Model | 0.03 | 0.09 |
| **60-69** |  |  |
| Ensemble of Age-Group Models | 0.32 | 0.54 |
| Independent Age-Group Models | 0.32 | 0.44 |
| Single Model | 0.44 | 0.24 |
| **70 Plus** |  |  |
| Ensemble of Age-Group Models | 0.38 | 0.65 |
| Independent Age-Group Models | 0.35 | 0.58 |
| Single Model | 0.12 | 0.38 |

### 3.3. Simulation of scenarios with the hybrid model

Five different scenarios are simulated by the hybrid model, and each time two related scenarios are compared. The Five scenarios are summarized in Table 3. In all these scenarios, the person whose variables are simulated is, at the start, a 40-year-old man, with a start weight of 90 kilos, 36% fat, and a height of 185 cm.

**Table 3:** Risk factors for the 5 different scenarios.

|  | Weight<br>loss | Weight<br>gain | Blood<br>pressure<br>medication | Diabetes | Smoking | Atrial<br>fibrillation | Gender |
| --- | --- | --- | --- | --- | --- | --- | --- |
| Scenario 1 |  | X |  | X |  |  | Male |
| Scenario 2 | X |  |  |  |  |  | Male |
| Scenario 3 |  | X | X |  |  |  | Male |
| Scenario 4 |  | X |  | X | X | X | Male |
| Scenario 5 | X |  | X |  |  |  | Male |

#### 3.3.1 Scenario 1 and 2

In scenario 1 (Figure 9A), a 40-year-old person that eats approximately 400 extra kcal/day for 40 years, goes from a BMI of around 28, indicating a slight overweight, to a BMI of almost 40, indicating obesity. In contrast, in scenario 2 (Figure 9B), the same person instead decreases their energy intake with 100 kcal/day and therefore has a BMI of 22 after 40 years. When zooming in and comparing the meal response for scenario 1 at age 40 and 80 (Figure 9A), one can see that for example the plasma glucose levels are higher both at start and at its peak, at age 80 compared with age 40. Insulin, on the other hand, is lower at age 80, due to the beta cell collapse, meaning that the person has gone from pre-diabetes to fully developed type 2 diabetes. Another difference that can be seen is that the glucose uptake in muscle tissue has decreased at age 80, but not in the expanded adipose tissue, which is another hallmark of diabetes. When looking at the risks for these two scenarios (Figure 9C), the risk increases slightly more for scenario 1, until age 50, when the diabetes kicks in, and the risk increases prominently. Note that the discrete nature of this jump probably is over-exaggerated, and due to the fact that the risk model is based on discrete diagnosis variable, and not on the more continuously changing physiological variables.

**Figure 9:**
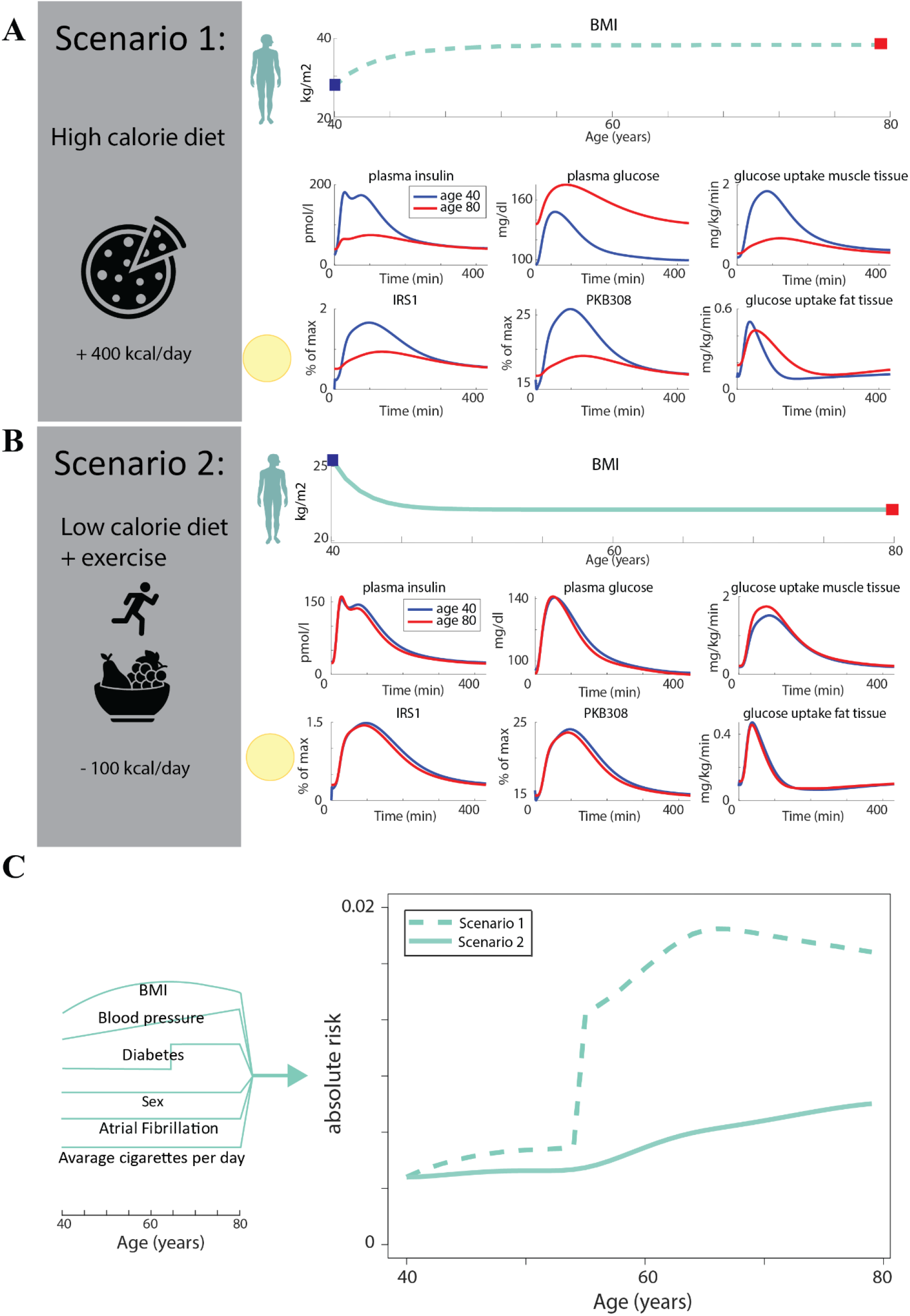
Simulations and comparisons of scenario 1 and 2. A) Scenario 1: A high-calorie diet, with a daily increase of 400 kcal, and a lack of exercise, leads to an increase of BMI over 40 years. The short-term dynamics at age 80 are here different from the ones at age 40 – plasma insulin has decreased, plasma glucose has increased to diabetic levels, glucose uptake in muscle tissue has decreased, glucose uptake in fat tissue is at the same total amount, but done over a period of time, and insulin-induced signaling IRS1 and PKB308 in isolated fat cells taken from the patient at these two time-points have decreased. B) Scenario 2: A low-calorie diet, decreasing the amount of food per day by 100 kcal, while also increasing the amount of exercise, which leads to a decrease in BMI over the 40 years. The short-term dynamics at age 40, before the weight decrease started, and at age 80, are more or less the same at both the organ/tissue and at the cell level. C) The risk of having a stroke as a function of age for the two scenarios. The risk is constantly higher for scenario 2.

**Figure 10:**
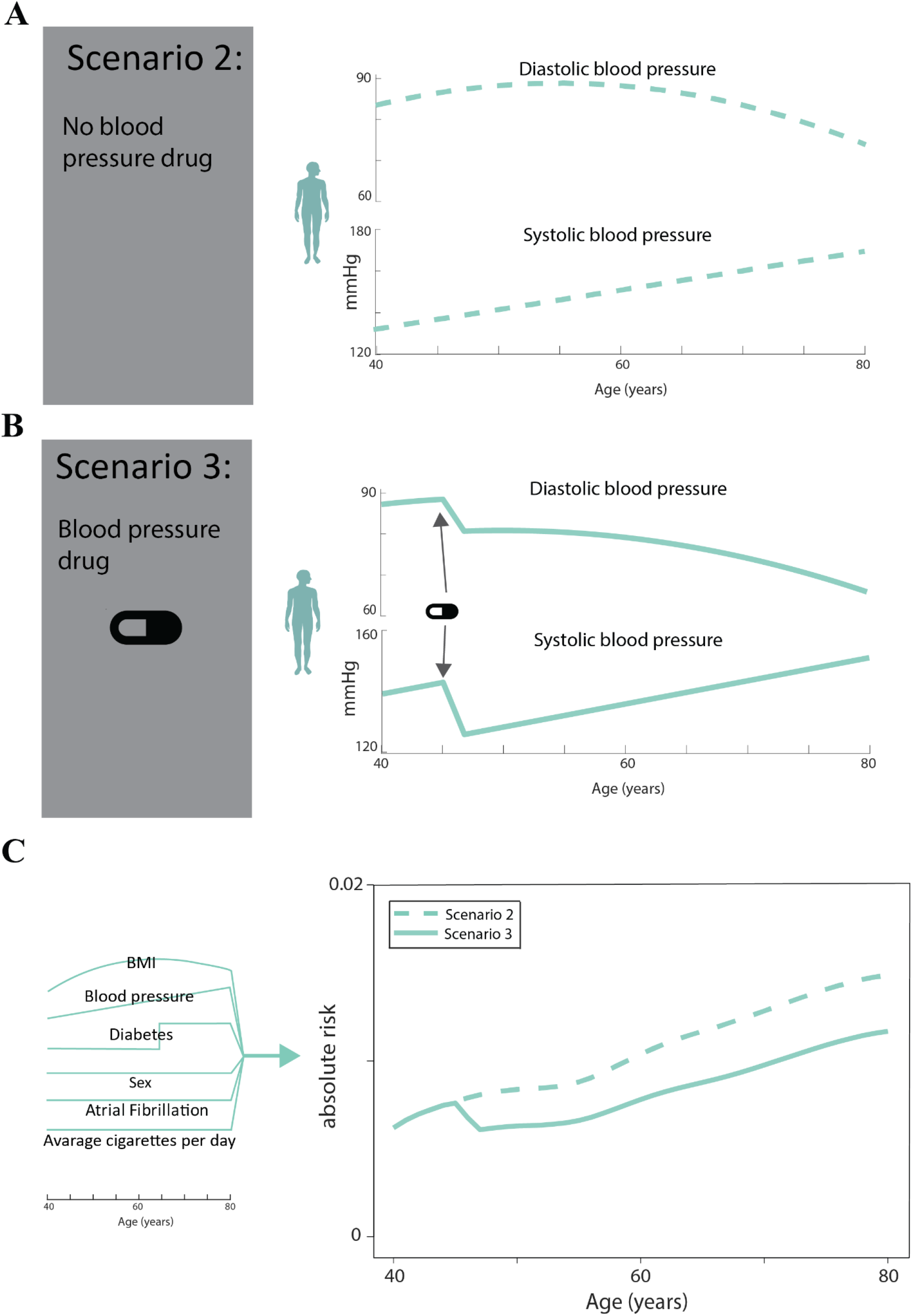
Simulations of scenario 2 and 3 compared. A) Scenario 2: No blood pressure drug is taken. B) Scenario 3: Blood pressure increases with age, and at age 50, the person starts taking a blood pressure drug. C) The risk of having a stroke as a function of age for the two scenarios. The risk is constantly higher for scenario 2, because of the blood pressure medication has taken in scenario 3.

**Figure 11:**
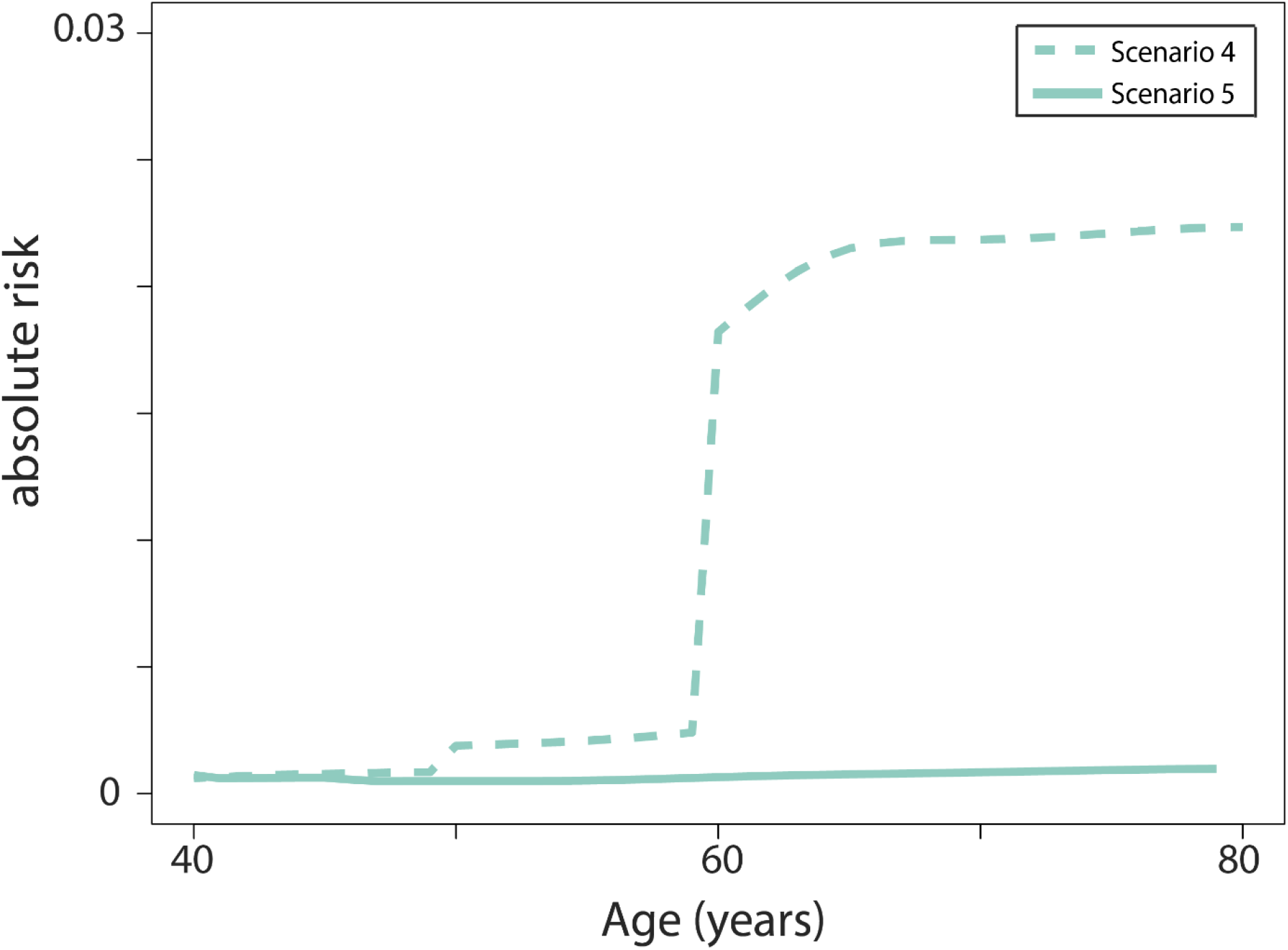
Risk scores for scenario 4 and 5, the worst- and best-case scenarios. In Scenario 4, a weight gain is compounded by continued smoking, diabetes, atrial fibrillation, and the omission of blood pressure medications. In Scenario 5, weight loss and quitting smoking is combined with blood pressure medication.

#### 3.3.2 Scenario 2 and 3

In scenario 3, the person starts taking a blood pressure medication at age 45, consequently decreasing both the SBP and DBP (Figure 10B). This scenario is compared with scenario 2 (Figure 10A), where no medication is taken, and the SBP thus consistently increases with time. Note, however, that the DBP increases at first, and then starts to decrease around age 50. All other risk factors are kept the same for these two scenarios. When comparing the risks for the two scenarios (Figure 10C), one can see a drop in the stroke risk for scenario 3 at the same time that the blood pressure medication is introduced. Note that this reduced risk is maintained throughout the remaining 30 years of simulated time.

#### 3.3.3 Scenario 4 and 5

Let us, finally, compare two more extreme scenarios, to see how much of a difference the lifestyle really can make (Fig 11). In the best-case scenario, the 40-year old reduces weight (as in scenario 1), stops smoking, and also takes a blood pressure medication. Just as in scenario 1, the risk is consistently low, throughout the 40 years of simulated time. In contrast, in scenario 5, the patient gains weight and does not take blood pressure medications, just as in scenario 2. However, here these outcomes are further compounded by the advent of diabetes and atrial fibrillation and continued smoking. As can be seen, the difference in absolute risk is more than 10 times higher in the worst-case scenario, compared to the best case. This shows how big of a difference lifestyle changes can make, and could be the basis of a new digital twin technology, useful in preventive healthcare.

### 3.4 Health informatics harmonization, to help bring the hybrid model into the clinic

If the digital twin models should become a ready-to-use eHealth technology, additional work needs to be done, for example concerning health informatics and end-usage design. There is some, but not much, previous work that focus on data management concerning digital twins (36). Some challenges mentioned include data heterogeneity, the potentially large size of data, the fact that the data is dynamic, i.e. changes over time, and the related data mining problems that these challenges give rise to (36).

Here we focus on the data heterogeneity and the need for semantic harmonization and integration. In the healthcare domain, future digital twins will need to access large amounts of complex heterogeneous data (e.g. omics data, EHR data, wearables data, etc.) which are heterogeneously structured and represented using a wide variety of formats.

There are multiple standards to enable data sharing among systems such as the EHR data exchange standards ISO 13606 (37), HL7 FHIR (38), and openEHR (39) as well as the OHDSI OMOP (40) and i2b2 (41) approaches, which are more oriented to clinical research. Thus, one of the main challenges is to deliver methods that enable the meaningful transformation of data across the existing heterogeneous representation formats. Among others, such methods will need to support the building of federated architectures for digital twins (42,43).

Within the Precise4Q project, where these digital twins for stroke have been developed, we have implemented a semantic-driven framework for semantically harmonizing and integrating data across systems, which facilitates data analysis and exploitation. A data transformation pipeline semantically harmonizes and standardizes the data input to the digital twins. The core of the framework is an ontology-based data model, which uses a top-level ontology to standardize data modelling and ensure interoperability among different ontologies, and the SNOMED CT terminology is used as a reference ontology to represent the clinical concepts.

Data from the different sources are usually provided as comma-separated values (CSV). We have therefore implemented a data transformation pipeline that takes CSV data as input, transforms it into RDF triples (44) according to the ontology data model, and stores it in a graph-database. Finally, data can be accessed by ML models by using a REST API. Figure 12 depicts the graphical representation of harmonized data from two datasets with patient examination and diagnostic data used in the models herein. Dashed grey squares are SNOMED CT concepts that univocally identify the data elements (e.g. BMI, SPB, Atrial fibrillation, etc.). These variables are related according to the mentioned ontology-based data model. Each data element is always represented together with its contextual information given by the specific process used to acquire the data (e.g. blood pressure taking process at some specific time point). This graph-based structure allows for further exploitation of biomedical knowledge and relations within the heterogenous data, for example for predicting new relations between terms such as genes, diseases, drugs, or symptoms (45,46).

**Figure 12:**
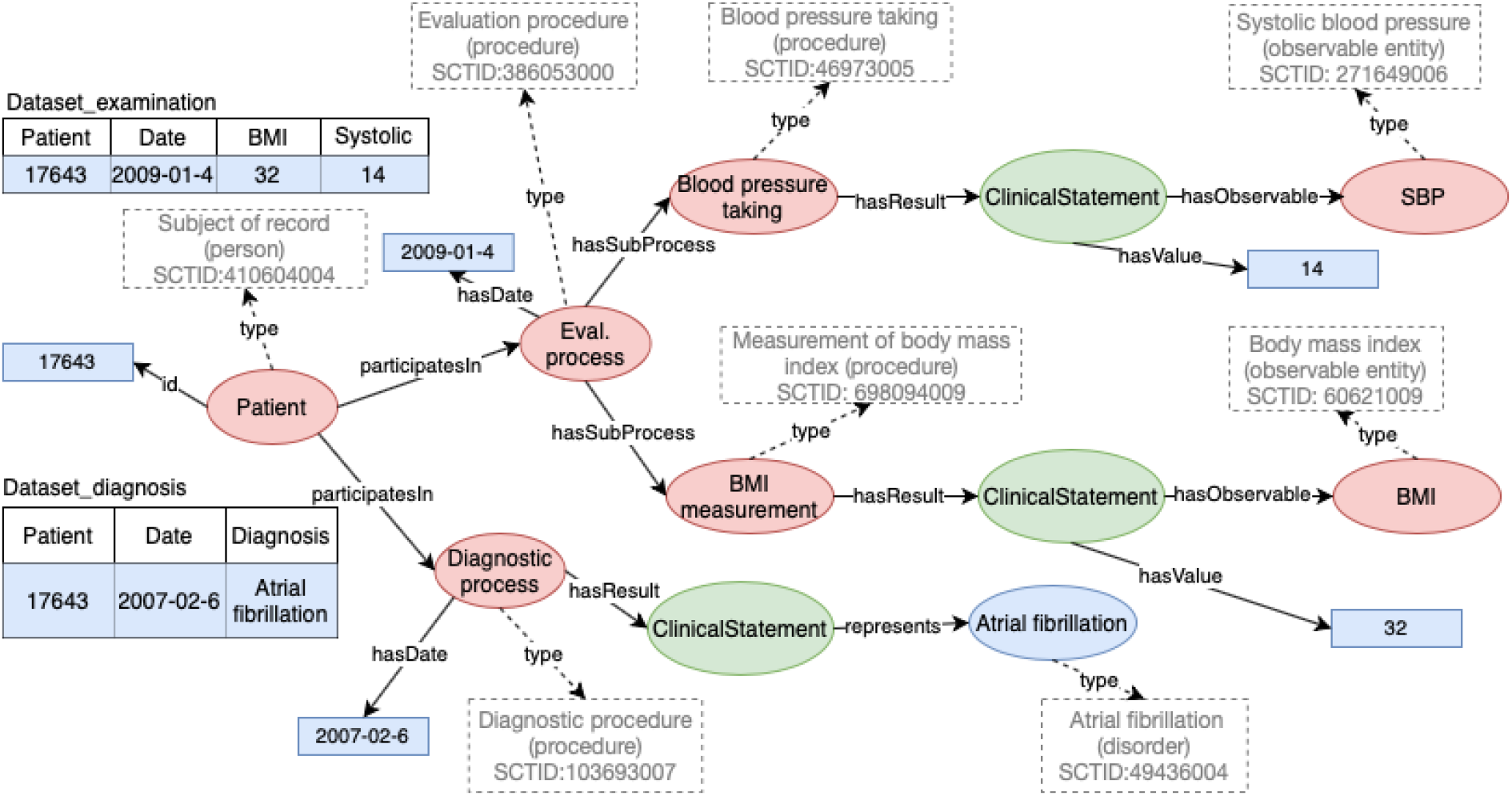
Simplified representation of the result of transforming csv data from two datasets into RDF triples according to the ontology model. Data is shown in blue. SCT concepts are represented as dashed grey squares

## 4. Discussion

### 4.1. Summary

Herein, we have presented the basis for a new digital twin technology, which uses a hybrid combination of a machine learning model with a multi-level, multi-timescale, mechanistic model. This hybrid model can show both the evolution of stroke risk and simulations of physiological variables, such as the progression of diabetes, weight, and blood pressure. The new hybrid model is based on a combination of existing and new models. The multi-level and multi-timescale model describes weight changes and glucose homeostasis (28,29). One of the new models (Figure 4) describes blood pressure as a function of age (Figure 5-7) and medications (Figure 7), and can both agree with estimation data (Figure 5,7), and correctly predict validation data (Figure 6). The second new model is a modification of the stroke risk model from (31), using a new ensemble approach (Figure 8). This ensemble approach was introduced to smooth out discontinuities, when moving from one age group model to another, and we show that the performance of the new model is as good as the previous model (31), both regarding discrimination (Table 1) and calibration (Table 2). We illustrate how the hybrid model can be used, by simulating and comparing different lifestyle scenarios: for example comparing the health trajectory for an individual who gains weight and develops diabetes (Figure 9, scenario 1) with that of an individual who loses weight and thereby improves their glucose meal response (Figure 9-10, scenario 2); other scenarios considered include an individual who gains weight but take blood pressure medication (Figure 10, scenario 3), an individual who smokes and develops atrial fibrillation (Figure 11, scenario 4), and an individual who loses weight and takes blood pressure medication (Figure 11, scenario 5). Finally, we also discuss and exemplify how health informatics solutions could be used to facilitate the usage of our hybrid model, both in the clinic and at home.

### 4.2. Strengths of our new models

#### New blood pressure model

The blood pressure model is new because it can describe both how blood pressure increases with age and how medication lowers the blood pressure. In the literature, there exist several models of the regulation of blood pressure, but only a few of them focus on long-term regulation. Guyton’s models created the groundwork for modeling long-term regulation of blood pressure based on simulations and animal data, including detailed descriptions of the salt-fluid balance (47,48). After Guyton’s model, many similar models have been developed (49), along with short-term Windkessel- (50) and time-varying elastance (51)-based models with more focus on the arterial system and the heart (52,53). For instance, Maksuti et al used a Windkessel-based model to describe blood pressure changes in aging using some of the Framingham data (54). However, the Maksuti model only describes blood pressure changes during normal aging, thus leaving out the three other groups in the Framingham data (Figure 5-6), and did not include effects of blood pressure-lowering drugs, as in the model presented herein (Figure 7). Apart from these models, only a few models describe the dynamics of long-term changes in systolic as well as diastolic blood pressure, such as aging and effects of anti-hypertensive drugs (49,54). Additionally, many of the previous models are not, or are only sparsely, based on human data, do not consider parameter estimation or validation with independent data, and/or do not combine several effects, such as both drugs and normal and hypertensive aging (49,54).

#### New ensemble-based risk model

The ensemble-based machine learning model deals with many of the most important risk factors, does so in an age-dependent way, while avoiding discontinuities, when moving from one age group to another. The model enables different risk factors to contribute to the risk scores in different amounts at different stages of life by having different models for different age groups. This is important, because the same risk factors will have different contributions in different stages of life, meaning that it is not sufficient to have age as an independent risk factor, along with the other ones (31). Moreover, the model generates smooths changes in risk score when entering a new age group by using a combined risk score for all age groups. Without this combination, an individual that ages and moves between age groups may display an unrealistic jump, between two risks. For example, a 49-year-old might see a decrease in risk score when they turn 50, even though none of their risk factors have changed. Another reason why this new ensemble-based approach is valuable is that there can be a difference between a person’s chronological age, which would be used to select the appropriate model to predict a person’s age, and their biological age. A number of studies have shown that stroke patients tend to have higher biological ages compared to non-stroke patients with the same chronological age (55). For this reason, it may be beneficial to down-play the importance of chronological age, and focus more on the other risk factors, which are a reflection of the biological age. This down-playing of chronological age appears since we use a weighted combination age-specific sub-models, thereby smoothly integrating information from different age cohorts. In these sub-models, age is less important, compared to in all-age models, where age is a more dominant risk factor.

#### The new hybrid model

The combined hybrid model is the first of its kind for stroke care, since it both can predict the risk of stroke, as well as predict the evolution of risk factors and other physiological variables, ranging from intracellular insulin signaling to organ fluxes, and whole-body variables. This combined ability is possible from our sequential hybrid modelling approach, where the mechanistic model is simulated first, and where that serves as an input to the stroke risk model. This combined ability is important in a clinical context, since one then needs to both see how the risk is expected to change as a function of lifestyle, and understand why these changes are happening, i.e. see the underlying physiological changes (Figure 1). Furthermore, it is also important to be able to simulate on a both long and short timescale, since it is only via short-term simulations (days-months) that one can compare predictions with observed changes and thus gain more faith in the models, and since it is only via long-term simulations (years-decades) that risks are substantially altered. Finally, as mentioned in the introduction, there are some, but not many, hybrid models in biomedicine already available (20–27), but no other model exists for multi-scale simulations of stroke risk.

### 4.3. Limitations of our models

#### Multi-level multi-timescale model

The multi-level, multi-timescale mechanistic model used herein constitutes a first step towards a model of stroke progression, but it has as several crucial limitations. Firstly, the weight-driven insulin resistance is constructed in a top-down fashion. The top-level is the weight-change, which is altered by changes to the balance between energy intake and consumption. These top-level weight-changes then affect tissue and cellular processes. However, the short-term dynamics on the lower levels does not propagate up to the top-level. This is a limitation, since long-term changes are nothing else than the cumulative impact of short-term changes, and this means that short-term changes seen during a few days do not impact the long-term predictions. Second, some of the organs and tissues represented are described by highly simplified sub-models. Therefore, in such organs, e.g. muscle, liver and brain, only a few variables can be simulated, such as glucose uptake. There are more detailed models for these organs (56–58), but these models have not yet been integrated into our multi-level model. A third limitation is the lack of other processes related to stroke represented in the mode. Now, the model focuses on the development of insulin resistance and diabetes, which are important in the early etiology of the metabolic syndrome. Future versions of our digital twin models should include mechanisms such as the development of arterial fibrillation, atherosclerosis, and thrombosis.

#### New blood pressure model

While our new blood pressure model is the first to both describe changes with age and medications, and while it is able to both describe estimation data and correctly predict independent validation data, it is still a simple, isolated, and phenomenological model. First, the new blood pressure model is not based on mechanistic Windkessel models, and can thus not simulate short-term dynamics, such as the blood pressure propagation during a single heartbeat. Second, for the same reason, our model is phenomenological, and predicts the long-term changes in systolic and diastolic blood pressure, without predicting the underlying changes in the mechanisms that are involved in regulating the blood pressure. For instance, the model does not include changes to arterial properties, such as stiffness and elastance, or changes to kidneys, etc. The reason for this limitation is that the required data for such a model - e.g. 4D-flow MRI data collected over long-term longitudinal studies - does not yet exist. Finally, the blood pressure model is not connected to the multi-level, multi-timescale model for weight and glucose homeostasis, even though blood flow has a mutual interplay with e.g. glucose uptake and diabetes (28).

#### New ensemble-based risk model

The risk model presented herein includes 7 of the most important risk factors for stroke (Figure 4), but there are several other risk factors that are not included. These risk factors include other heart diseases, birth control pills, previous strokes or TIAs, high red blood count, high blood cholesterol and lipids, alcohol and drug use, cardiac structural abnormalities, and social and economic factors. All these risk factors are manageable to some extent at least, and not including them in a risk model used in the clinic could mean that important possible changes to reduce the risk are overlooked. To include more risk factors would, however, also mean that missing values need to be addressed in some way, both during training and during usage of the risk model. The risk factors currently included are all easy to know or measure, and are measured in all known databases, and this is the case also for several other risk factors not included. Such variables would be straightforward to add to our model, and missing values could be handled by using some imputation method. In contrast, some other risk factors are measured only rarely, and are thus not included in all large databases. For instance, cardiac structural abnormalities are diagnosed using echocardiography, which is done in the clinic only if there is a strong suspicion of disease, and not a part the Framingham study, used for our risk models. Addition of such variables would thus require the usage of other databases, combine with usage of a transfer learning approach to merge the insights from analysis of such different databases (59).

#### The new hybrid model

Our hybrid risk model for stroke is the first of its kind, and is to be considered a first step, with many future improvements possible. The limitations of the different sub-models of course carry over to the combined hybrid model as well. Apart from this, there are several hybrid approaches that are not utilized in our model. The approach that we do use, sequential hybrid modelling, is the simplest one, since you can simulate one model first, and then use that as an input to the second model. A more intricate form of hybrid modelling is called blended modelling, where the different models interact with each other, and thus must be analyzed together. One way to do such blended modelling, would be to use nonlinear mixed-effects modelling (60,61), which is a way of bringing in statistical properties, regarding e.g. the distribution of parameters across a population, into the mechanistic simulations. Another way to do such blended modelling would be to have a sub-model to the big multi-level model that is described using a phenomenological machine-learning model. Finally, another important sequential hybrid combination not pursued herein, would be to use a machine-learning or bioinformatics network model as an input to the ultimate risk model. Such a model could e.g. be a polygenic risk score model, which takes all the genes and calculate a smaller set of risk factors, which then goes into the ultimate risk model. These different future possibilities are reviewed for the context of stroke in (10).

### 4.4. Future steps towards potential clinical implementation

The hybrid model presented herein opens up for a future, fully realized, implementation of P4 medicine in health care, i.e. Predictive, Preventive, Personalized and Participatory healthcare (Figure 1). The Predictive part comes through the predictive simulations and risk scores (Figure 9-11, the scenarios); the Preventive part comes from using the predictions in e.g. the preventive health dialogue; Personalization comes through the fact that the models are personalized; and finally, the Participatory part comes from the fact that the simulation empowers patients to make informed decisions about their health behaviors, lifestyle choices, and interventions, not just in connection to the Health Conversation, but continuously throughout life. This last potential was also mentioned in the introduction (Figure 1). Short-term simulations can be compared with data and can thus be used to examine and verify the short-term effects (e.g. glucose meal response) of the long-term changes (e.g. weight changes). This can be done whenever the patient so decides, i.e. after a year, half a year, or even a day. By improving patients’ understanding of how different health and lifestyle factors and changes will impact their stroke risk, the digital twin can serve as a valuable tool for patient education and as a conversation aid during the clinical encounter. Being able to simulate how different types of interventions or lack thereof may reduce or increase the risk of suffering a stroke, enables patients to make informed decision about their health at eye level with healthcare professionals. Moreover, if predictions are consistent with the observed changes, this can increase patients’ trust in the model, including for the long-term predictions, and as a result increase their motivation and compliance to lifestyle changes.

Despite the great potential of predictive health information to empower individuals, it is important to consider some of the pitfalls and potential challenges patients may face in making effective use of this information. Even though the digital twin can foster awareness of much needed lifestyle changes, it does not help patients to overcome the environmental or social factors that may prevent them from adopting a healthier lifestyle. It is also important to note that not all patients are willing and capable to take on such an active role in managing their own health and may feel burdened by the responsibility of having to do so. This is where healthcare professionals are called upon to support patients in identifying not only the right type of lifestyle intervention but also how it can be achieved.

Finally, when bringing the herein presented hybrid model all the way to an implemented eHealth technology, there are several additional steps needed. For instance, data from different sources need to be integrated and harmonized in a secure way. User interfaces need to be co-designed with end-users, including both patients, clinicians, and other persons that may be involved, such as personal trainers, coaches, relatives, and friends.

