## Supplementary for "Digital twins and hybrid modelling for simulation of physiological variables and stroke risk"

### 1 System of ordinary different equations

The ODEs in the model are given in Eq. (3).

$$
\begin{aligned}
d/dt(Gly) &= v1/rhog \\
d/dt(ECF) &= v2 \\
d/dt(F) &= (1 - v3) \cdot (k1 - v1)/rhof \\
d/dt(L) &= v3 \cdot (k1 - v1)/rhol \\
d/dt(AT) &= v4 \\
d/dt(dosage) &= 0 \\
d/dt(RE) &= 0 \\
d/dt(G\_p) &= (EGP + Ra - E - U\_ii - k\_1 \cdot G\_p + k\_2 \cdot G\_t) \cdot tconv \\
d/dt(G\_t) &= ((-U\_id) + k\_1 \cdot G\_p - k\_2 \cdot G\_t) \cdot tconv \\
d/dt(I\_l) &= ((-m\_1 \cdot I\_l) - m\_3 \cdot I\_l + m\_2 \cdot I\_p + S) \cdot tconv \\
d/dt(I\_p) &= ((-m\_2 \cdot I\_p) - m\_4 \cdot I\_p + m\_1 \cdot I\_l) \cdot tconv \\
d/dt(Q\_sto1) &= (-k\_gri \cdot Q\_sto1) \cdot tconv \\
d/dt(Q\_sto2) &= ((-k\_empt \cdot Q\_sto2) + k\_gri \cdot Q\_sto1) \cdot tconv \\
d/dt(Q\_gut) &= ((-k\_abs \cdot Q\_gut) + k\_empt \cdot Q\_sto2) \cdot tconv \\
d/dt(I\_1) &= -k\_i \cdot (I\_1 - I) \cdot tconv \\
d/dt(I\_d) &= -k\_i \cdot (I\_d - I\_1) \cdot tconv \\
d/dt(I\_po) &= ((-gamma \cdot I\_po) + S\_po) \cdot f\_IR\_Ins \cdot tconv \\
d/dt(Y) &= (-alpha \cdot (Y - beta \cdot (G - G\_b))) \cdot tconv \\
d/dt(INS\_f) &= ((-p\_2U \cdot INS\_f) + p\_2U \cdot (I - I\_b)) \cdot tconv \\
d/dt(INS) &= (V\_1 - V\_2) \cdot tconv \\
d/dt(betamass) &= (-d0 + r1 \cdot G - r2 \cdot G \cdot G) \cdot betamass/tconv \\
d/dt(IR) &= (-v1a - v1basal + v1r + v1g) \cdot tconv \\
d/dt(IRp) &= (v1basal + v1c - v1d - v1g) \cdot tconv \\
d/dt(IRins) &= (v1a - v1c) \cdot tconv \\
d/dt(IRip) &= (v1d - v1e) \cdot tconv \\
d/dt(IRi) &= (v1e - v1r) \cdot tconv \\
d/dt(IRS1) &= (v2b + v2g - v2a - v2basal) \cdot tconv \\
d/dt(IRS1p) &= (v2a + v2d - v2b - v2c) \cdot tconv \\
d/dt(IRS1p307) &= (v2c - v2d - v2f) \cdot tconv \\
d/dt(IRS1307) &= (v2basal + v2f - v2g) \cdot tconv \\
d/dt(X) &= (v3b - v3a) \cdot tconv \\
d/dt(Xp) &= (v3a - v3b) \cdot tconv
\end{aligned}
\tag{1}
$$

$$
\begin{aligned}
d/dt(PKB) &= (-v4a + v4b + v4h) \cdot tconv \\
d/dt(PKB308p) &= (v4a - v4b - v4c) \cdot tconv \\
d/dt(PKB473p) &= (-v4e + v4f - v4h) \cdot tconv \\
d/dt(PKB308p473p) &= (v4c + v4e - v4f) \cdot tconv \\
d/dt(mTORC1) &= (v5b - v5a) \cdot tconv \\
d/dt(mTORC1a) &= (v5a - v5b) \cdot tconv \\
d/dt(mTORC2) &= (-v5c + v5d) \cdot tconv \\
d/dt(mTORC2a) &= (v5c - v5d) \cdot tconv \\
d/dt(AS160) &= (v6b1 - v6f1) \cdot tconv \\
d/dt(AS160p) &= (v6f1 - v6b1) \cdot tconv \\
d/dt(GLUT4m) &= (v7f - v7b) \cdot tconv \\
d/dt(GLUT4) &= (-v7f + v7b) \cdot tconv \\
d/dt(S6K) &= (v9b1 - v9f1) \cdot tconv \\
d/dt(S6Kp) &= (v9f1 - v9b1) \cdot tconv \\
d/dt(S6) &= (v9b2 - v9f2) \cdot tconv \\
d/dt(S6p) &= (v9f2 - v9b2) \cdot tconv \\
d/dt(Glu\_in) &= (p1 \cdot (V\_in - V\_out) - V\_G6P) \cdot tconv \\
d/dt(G6P) &= (V\_G6P - V\_met) \cdot tconv
\end{aligned} \tag{2}
$$

### 2 Initial values

The initial values of the model states are given in Eq. (4) and **??**. Eq. (4) is given below.

$$
\begin{aligned}
Gly(0) &= 1.8500 \\
ECF(0) &= 1.4805 \cdot 10^{1} \\
F(0) &= 2.3400 \cdot 10^{1} \\
L(0) &= 4.4950 \cdot 10^{1} \\
AT(0) &= 1.5963E{-}28 \\
dosage(0) &= 0.0000 \\
RE(0) &= 0.0000 \\
G\_p(0) &= 1.7800 \cdot 10^{2} \\
G\_t(0) &= 1.3000 \cdot 10^{2} \\
I\_l(0) &= 4.5000 \\
I\_p(0) &= 1.2500 \\
Q\_sto1(0) &= 7.8000 \cdot 10^{4} \\
Q\_sto2(0) &= 0.0000 \\
Q\_gut(0) &= 0.0000 \\
I\_1(0) &= 2.5000 \cdot 10^{1} \\
I\_d(0) &= 2.5000 \cdot 10^{1} \\
I\_po(0) &= 3.6000 \\
Y(0) &= 0.0000 \\
INS\_f(0) &= 0.0000 \\
INS(0) &= 0.0000 \\
betamass(0) &= 2.1020 \cdot 10^{2} \\
IR(0) &= 9.8951 \cdot 10^{1} \\
IRp(0) &= 2.0651 \cdot 10^{-3} \\
IRins(0) &= 9.1254 \cdot 10^{-1} \\
IRip(0) &= 1.7520 \cdot 10^{-2} \\
IRi(0) &= 1.1706 \cdot 10^{-1} \\
IRS1(0) &= 8.6327 \cdot 10^{1} \\
IRS1p(0) &= 1.4248 \cdot 10^{-3} \\
IRS1p307(0) &= 6.4102 \cdot 10^{-4} \\
IRS1307(0) &= 1.3671 \cdot 10^{1} \\
X(0) &= 9.9998 \cdot 10^{1} \\
Xp(0) &= 1.9871 \cdot 10^{-3}
\end{aligned}
\tag{3}
$$

$$
\begin{aligned}
PKB(0) &= 6.6321 \cdot 10^{1} \\
PKB308p(0) &= 1.5444 \cdot 10^{1} \\
PKB473p(0) &= 1.8164 \cdot 10^{1} \\
PKB308p473p(0) &= 7.1290 \cdot 10^{-2} \\
mTORC1(0) &= 9.6196 \cdot 10^{1} \\
mTORC1a(0) &= 3.8041 \\
mTORC2(0) &= 9.9859 \cdot 10^{1} \\
mTORC2a(0) &= 1.4151 \cdot 10^{-1} \\
AS160(0) &= 8.7467 \cdot 10^{1} \\
AS160p(0) &= 1.2533 \cdot 10^{1} \\
GLUT4m(0) &= 2.1845 \cdot 10^{1} \\
GLUT4(0) &= 7.8155 \cdot 10^{1} \\
S6K(0) &= 9.9790 \cdot 10^{1} \\
S6Kp(0) &= 2.1007 \cdot 10^{-1} \\
S6(0) &= 9.7794 \cdot 10^{1} \\
S6p(0) &= 2.2063 \\
Glu\_in(0) &= 2.4710 \\
G6P(0) &= 4.2738 \cdot 10^{-3}
\end{aligned}
\tag{4}
$$

### 3 Parameter values

The parameter values of the model states are given in Table 1-4.

| Parameter | Value |
|---|---|
| $diabetes$ | 1.0000 |
| $k1a$ | $6.3314 \cdot 10^{-1}$ |
| $k1basal$ | $3.3134 \cdot 10^{-2}$ |
| $k1c$ | $8.7680 \cdot 10^{-1}$ |
| $k1d$ | $3.1012 \cdot 10^{1}$ |
| $k1f$ | $1.8396 \cdot 10^{3}$ |
| $k1g$ | $1.9441 \cdot 10^{3}$ |
| $k1r$ | $5.4706 \cdot 10^{-1}$ |
| $k2a$ | 3.2273 |
| $k2c$ | $5.7588 \cdot 10^{3}$ |
| $k2basal$ | $4.2277 \cdot 10^{-2}$ |
| $k2b$ | $3.4243 \cdot 10^{3}$ |
| $k2d$ | $2.8075 \cdot 10^{2}$ |
| $k2f$ | 2.9131 |
| $k2g$ | $2.6709 \cdot 10^{-1}$ |
| $k3a$ | $1.3773 \cdot 10^{-3}$ |
| $k3b$ | $9.8756 \cdot 10^{-2}$ |
| $k4a$ | $5.7902 \cdot 10^{3}$ |
| $k4b$ | $3.4797 \cdot 10^{1}$ |
| $k4c$ | 4.4558 |
| $k4e$ | $4.2840 \cdot 10^{1}$ |
| $k4f$ | $1.4360 \cdot 10^{2}$ |
| $k4h$ | $5.3614 \cdot 10^{-1}$ |
| $k5a1$ | 1.8423 |
| $k5a2$ | $5.5064 \cdot 10^{-2}$ |
| $k5b$ | $2.4826 \cdot 10^{1}$ |
| $k5d$ | 1.0601 |
| $km5$ | 2.6499 |
| $k5c$ | $8.5751 \cdot 10^{-2}$ |
| $k6f1$ | 2.6517 |
| $k6f2$ | $3.6935 \cdot 10^{1}$ |

Table 1: **Parameter values.**

| | |
|---|---|
| $km6$ | $3.0542 \cdot 10^{1}$ |
| $n6$ | $2.1371$ |
| $k6b$ | $6.5184 \cdot 10^{1}$ |
| $k7f$ | $5.0983 \cdot 10^{1}$ |
| $k7b$ | $2.2860 \cdot 10^{3}$ |
| $k8$ | $7.2424 \cdot 10^{2}$ |
| $glut1$ | $7.0422 \cdot 10^{3}$ |
| $k9f1$ | $1.2981 \cdot 10^{-1}$ |
| $k9b1$ | $4.4409 \cdot 10^{-2}$ |
| $k9f2$ | $3.3289$ |
| $k9b2$ | $3.0997 \cdot 10^{1}$ |
| $km9$ | $5.8727 \cdot 10^{3}$ |
| $n9$ | $9.8547 \cdot 10^{-1}$ |
| $kbf$ | $1.0000 \cdot 10^{-2}$ |
| $nC$ | $2.1000 \cdot 10^{-6}$ |
| $p1$ | $1.7891 \cdot 10^{-1}$ |
| $p2$ | $4.4782$ |
| $p3$ | $1.6088 \cdot 10^{-1}$ |
| $p4$ | $2.6271$ |
| $k\_gluin$ | $2.1361$ |
| $k\_G6P$ | $1.1496 \cdot 10^{4}$ |
| $V\_G6Pmax$ | $4.1020 \cdot 10^{2}$ |
| $V\_G$ | $1.8800$ |
| $k\_1$ | $6.5000 \cdot 10^{-2}$ |
| $k\_2$ | $7.9000 \cdot 10^{-2}$ |
| $G\_b$ | $9.5000 \cdot 10^{1}$ |
| $V\_I$ | $5.0000 \cdot 10^{-2}$ |
| $m\_1$ | $1.9000 \cdot 10^{-1}$ |
| $m\_2$ | $4.8400 \cdot 10^{-1}$ |
| $m\_4$ | $1.9400 \cdot 10^{-1}$ |
| $m\_5$ | $3.0400 \cdot 10^{-2}$ |
| $m\_6$ | $6.4710 \cdot 10^{-1}$ |
| $HE\_b$ | $6.0000 \cdot 10^{-1}$ |
| $I\_b$ | $2.5000 \cdot 10^{1}$ |
| $S\_b$ | $1.8000$ |
| $k\_max$ | $5.5800 \cdot 10^{-2}$ |
| $k\_min$ | $8.0000 \cdot 10^{-3}$ |
| $k\_abs$ | $5.7000 \cdot 10^{-2}$ |
| $k\_gri$ | $5.5800 \cdot 10^{-2}$ |
| $f$ | $9.0000 \cdot 10^{-1}$ |
| $b$ | $8.2000 \cdot 10^{-1}$ |
| $dd$ | $1.0000 \cdot 10^{-2}$ |

Table 2: **Parameter values.**

| | |
|---|---|
| $k\_p1$ | 2.7000 |
| $k\_p2$ | $2.1000 \cdot 10^{-3}$ |
| $k\_p3$ | $9.0000 \cdot 10^{-3}$ |
| $k\_p4$ | $6.1800 \cdot 10^{-2}$ |
| $k\_i$ | $7.9000 \cdot 10^{-3}$ |
| $p\_2U$ | $3.3100 \cdot 10^{-2}$ |
| $K$ | 2.3000 |
| $alpha$ | $5.0000 \cdot 10^{-2}$ |
| $beta$ | $1.1000 \cdot 10^{-1}$ |
| $gamma$ | $5.0000 \cdot 10^{-1}$ |
| $k\_e1$ | $5.0000 \cdot 10^{-4}$ |
| $k\_e2$ | $3.3900 \cdot 10^{2}$ |
| $D$ | $7.8000 \cdot 10^{4}$ |
| $INS\_offset$ | 7.0000 |
| $be$ | 3.0000 |
| $bradykinin$ | 1.0000 |
| $pf$ | $3.4000 \cdot 10^{1}$ |
| $bf\_b$ | 3.0000 |
| $INS\_b$ | 0.0000 |
| $p\_bf$ | 1.0000 |
| $U\_ii$ | $8.3087 \cdot 10^{-1}$ |
| $kI2$ | $4.2884 \cdot 10^{-2}$ |
| $kI1$ | $4.7612 \cdot 10^{-2}$ |
| $V\_m$ | $8.8133 \cdot 10^{-1}$ |
| $V\_mx$ | $4.0936 \cdot 10^{-2}$ |
| $K\_m$ | $4.7635 \cdot 10^{2}$ |
| $V\_l$ | 2.0047 |
| $V\_lx$ | $4.3933 \cdot 10^{-2}$ |
| $K\_l$ | $3.5485 \cdot 10^{2}$ |
| $BWinit$ | $9.0000 \cdot 10^{1}$ |
| $height$ | $1.8500 \cdot 10^{2}$ |
| $age$ | $5.0000 \cdot 10^{1}$ |
| $RMRinit$ | $7.1025 \cdot 10^{6}$ |
| $Ginit$ | 1.8500 |
| $ECFinit$ | $1.4805 \cdot 10^{1}$ |
| $Finit$ | $2.3400 \cdot 10^{1}$ |
| $Linit$ | $4.4950 \cdot 10^{1}$ |
| $ATinit$ | $1.0000 \cdot 10^{-1}$ |
| $EIrestriction2$ | $4.0000 \cdot 10^{2}$ |
| $alfa$ | 1.0000 |
| $fCIn$ | $4.0000 \cdot 10^{-1}$ |
| $PAE$ | 1.5000 |

Table 3: **Parameter values.**

| | |
|---|---|
| $rhol$ | $7.6000 \cdot 10^{6}$ |
| $rhof$ | $3.9500 \cdot 10^{7}$ |
| $gf$ | $1.3000 \cdot 10^{4}$ |
| $gl$ | $9.2000 \cdot 10^{4}$ |
| $etal$ | $9.6000 \cdot 10^{5}$ |
| $etaf$ | $7.5000 \cdot 10^{5}$ |
| $rhog$ | $1.7600 \cdot 10^{7}$ |
| $Na$ | 3.2200 |
| $epNa$ | $3.0000 \cdot 10^{3}$ |
| $epCI$ | $4.0000 \cdot 10^{3}$ |
| $deltaNaDiet$ | 0.0000 |
| $bTEF$ | $1.0000 \cdot 10^{-1}$ |
| $bAT$ | $1.4000 \cdot 10^{-1}$ |
| $tAT$ | $1.4000 \cdot 10^{1}$ |
| $delta$ | 1.0000 |
| $k$ | 1.0000 |
| $m$ | 1.0000 |
| $lmax$ | 0.0000 |
| $IC\_50$ | 1.0000 |
| $h2$ | 1.0000 |
| $h1$ | 5.0000 |
| $dEI\_ss$ | $1.2265 \cdot 10^{3}$ |
| $t\_half$ | 1.4240 |
| $r1$ | $9.1000 \cdot 10^{-4}$ |
| $r2$ | $2.2000 \cdot 10^{-5}$ |
| $d0$ | 0.0000 |
| $ba$ | $4.8000 \cdot 10^{-3}$ |
| $k\_egp$ | $3.0000 \cdot 10^{-1}$ |
| $k\_uid$ | 1.0000 |
| $k\_s$ | $5.0000 \cdot 10^{-1}$ |

Table 4: **Parameter values.**

### 4 Variables

The variable equations are given in Eq. (37).

$$
\begin{aligned}
f\_IR\_EGP &= 1 + (k\_egp \cdot F/Finit) && (5)\\
f\_IR\_Ins &= 1 + (k\_s \cdot log(F/Finit)) && (6)\\
f\_IR\_CLGI &= 1 + (k\_uid \cdot log(F/Finit)) && (7)\\
CC &= 10.4 \cdot rhol/rhof && (8)\\
p &= CC/(CC + F) && (9)\\
EEinit &= PAE \cdot RMRinit && (10)\\
EIninit &= EEinit && (11)\\
CIninit &= fCIn \cdot EIninit && (12)\\
kg &= CIninit/(Ginit \cdot Ginit) && (13)\\
dEI\_init &= EIrestriction2 \cdot 4183 && (14)\\
EI\_vehicle &= EIninit + dEI\_init && (15)\\
EIn &= (1 - alfa) \cdot EIninit + alfa \cdot EI\_vehicle && (16)\\
CInvalue &= fCIn \cdot EI\_vehicle && (17)\\
CIn &= (1 - alfa) \cdot CIninit + alfa \cdot CInvalue && (18)\\
PAL &= PAE && (19)\\
BW &= (F + L + (1 + 2.7) \cdot Gly + ECF) && (20)\\
TEF &= bTEF \cdot (EIn - EIninit) && (21)\\
KK &= EEinit - (gf \cdot Finit + gl \cdot Linit + delta \cdot BWinit) && (22)\\
EE &= (-BW \cdot delta \cdot rhof \cdot rhol - KK \cdot rhof \cdot rhol - rhof \cdot rhol \cdot TEF - rhof \cdot && (23)\\
k1 &= EIn - EE && (24)\\
I &= I\_p/V\_I && (25)\\
G &= G\_p/V\_G && (26)\\
aa &= 5/2/(1 - b)/D && (27)\\
cc &= 5/2/dd/D && (28)\\
EGP &= (k\_p1 - (k\_p2 \cdot G\_p + k\_p3 \cdot I\_d + k\_p4 \cdot I\_po)) \cdot (f\_IR\_EGP) && (29)\\
E &= k\_e1 \cdot (G\_p - k\_e2) && (30)\\
S &= ba \cdot betamass \cdot gamma \cdot I\_po && (31)\\
HE &= (-m\_5 \cdot S) + m\_6 && (32)\\
m\_3 &= HE \cdot m\_1/(1 - HE) && (33)\\
Q\_sto &= Q\_sto1 + Q\_sto2 && (34)\\
Ra &= f \cdot k\_abs \cdot Q\_gut/BW && (35)\\
& && (36)
\end{aligned}
$$

$$
\begin{aligned}
k\_empt &= k\_min + (k\_max - k\_min)/2 \cdot (tanh(aa \cdot (Q\_sto - b \cdot D)) - tanh(cc \cdot (Q\_s \text{[illegible]}\\
bf\_f &= bradykinin \cdot (be + kbf \cdot (INS\_f + INS\_offset))\\
bfe\_f &= bf\_f\\
S\_po &= (Y + K \cdot (EGP + Ra - E - U\_ii - k\_1 \cdot G\_p + k\_2 \cdot G\_t)/V\_G + S\_b)\\
V\_mmax &= V\_m + V\_mx \cdot INS\\
V\_lmax &= V\_l + V\_lx \cdot INS\\
INS\_fe &= nC \cdot (k8 \cdot GLUT4m/pf + glut1/pf + bfe\_f)\\
V\_in &= p4 \cdot G\_t \cdot INS\_fe\\
V\_out &= p3 \cdot Glu\_in\\
U\_idm &= (L/Linit) \cdot V\_mmax \cdot G\_t/(K\_m + G\_t)/f\_IR\_CLGI\\
U\_idl &= V\_lmax \cdot G\_t/(K\_l + G\_t)/f\_IR\_CLGI\\
U\_idf &= (F/Finit) \cdot (V\_in - V\_out)/f\_IR\_CLGI\\
U\_id &= (U\_idf + U\_idm + U\_idl)\\
U &= U\_id + U\_ii\\
measuredIRS1 &= IRS1p + IRS1p307\\
measuredPKB308 &= PKB308p + PKB308p473p\\
tconv &= 24 \cdot 60
\end{aligned}
\tag{37}
$$

### 5 Reactions

The reaction equations are given in Eq. (39).

$$
\begin{aligned}
v1 &= (CIn - kg \cdot (Gly)^2) \\
v2 &= 0 \\
v3 &= p \\
v4 &= (bAT \cdot (EIn - EIninit) - AT)/tAT \\
V\_1 &= kI1 \cdot (I - I\_b) \\
V\_2 &= kI2 \cdot INS \\
v1a &= IR \cdot k1a \cdot (INS\_f + 5) \cdot 1e - 3 \\
v1basal &= k1basal \cdot IR \\
v1c &= IRins \cdot k1c \\
v1d &= IRp \cdot k1d \\
v1e &= IRip \cdot k1f \cdot Xp \\
v1g &= IRp \cdot k1g \\
v1r &= IRi \cdot k1r \\
v2a &= IRS1 \cdot k2a \cdot IRip \\
v2b &= IRS1p \cdot k2b \\
v2c &= IRS1p \cdot k2c \cdot mTORC1a \cdot diabetes \\
v2d &= IRS1p307 \cdot k2d \\
v2f &= IRS1p307 \cdot k2f \\
v2basal &= (IRS1) \cdot k2basal \\
v2g &= IRS1307 \cdot k2g \\
v3a &= X \cdot k3a \cdot IRS1p \\
v3b &= Xp \cdot k3b \\
v5a &= mTORC1 \cdot (k5a1 \cdot PKB308p473p + k5a2 \cdot PKB308p) \\
v5b &= mTORC1a \cdot k5b \\
v5c &= mTORC2 \cdot k5c \cdot IRip \\
v5d &= k5d \cdot mTORC2a \\
v4a &= k4a \cdot PKB \cdot IRS1p \\
v4b &= k4b \cdot PKB308p \\
v4c &= k4c \cdot PKB308p \cdot mTORC2a \\
v4e &= k4e \cdot PKB473p \cdot IRS1p307 \\
v4f &= k4f \cdot PKB308p473p \\
v4h &= k4h \cdot PKB473p \\
v6f1 &= AS160 \cdot (k6f1 \cdot PKB308p473p + k6f2 \cdot PKB473p^n6/(km6^n6 + PKB473p^n6)) \\
v6b1 &= AS160p \cdot k6b
\end{aligned}
\tag{38}
$$

$$
\begin{aligned}
v7f &= GLUT4 \cdot k7f \cdot AS160p \\
v7b &= GLUT4m \cdot k7b \\
v9f1 &= S6K \cdot k9f1 \cdot mTORC1a^n9/(km9^n9 + mTORC1a^n9) \\
v9b1 &= S6Kp \cdot k9b1 \\
v9f2 &= S6 \cdot k9f2 \cdot S6Kp \\
v9b2 &= S6p \cdot k9b2 \\
V\_G6P &= V\_G6Pmax \cdot Glu\_in/(k\_gluin + Glu\_in) \cdot 1/(k\_G6P + G6P) \\
V\_met &= p2 \cdot G6P
\end{aligned}
\tag{39}
$$
